# Early-life nutrition interacts with developmental genes to shape the brain and sleep behavior in *Drosophila melanogaster*

**DOI:** 10.1101/2020.06.28.175356

**Authors:** Gonzalo H. Olivares, Franco Núñez-Villegas, Noemi Candia, Karen Oróstica, Franco Vega-Macaya, Nolberto Zúñiga, Cristian Molina, Trudy F. C. Mackay, Ricardo A. Verdugo, Patricio Olguín

## Abstract

The mechanisms by which the genotype interacts with nutrition during development to contribute to the variation of complex behaviors and brain morphology of adults are not well understood. Here we use the *Drosophila* Genetic Reference Panel to identify genes and pathways underlying these interactions in sleep behavior and mushroom body morphology. We show that early-life nutritional restriction has genotype-specific effects on variation in sleep behavior and brain morphology. We mapped genes associated with sleep sensitivity to early-life nutrition, which were enriched for protein-protein interactions responsible for translation, endocytosis regulation, ubiquitination, lipid metabolism, and neural development. By manipulating the expression of candidate genes in the mushroom bodies and all neurons, we confirm that genes regulating neural development, translation and insulin signaling contribute to the variable response of sleep and brain morphology to early-life nutrition. We show that the interaction between differential expression of candidate genes with nutritional restriction in early life resides in the mushroom bodies or other neurons, and that these effects are sex specific. Natural variation in genes that control the systemic response to nutrition and brain development and function interact with early-life nutrition in different types of neurons to contribute to the variation of brain morphology and adult sleep behavior.

## Introduction

Nutrition is an environmental factor that plays a crucial role in the maturation and functional development of the central nervous system ^1–3^. In mammals, including humans, severe prenatal malnutrition negatively impacts neural development and complex behaviors such as sleep, memory, and learning ^1, 4–7^. At the population level, adults that were exposed to hunger *in utero* have an increased risk to develop schizophrenia, affective disorders, addiction and decreased cognitive function ^8, 9^. The origin of these disorders may be associated with defects in early brain development ^10^.

Little is known about the mechanisms by which individual genotypes (G) respond differently to a nutritional environment (E) during development. We define this type of genotype by environment interaction (GEI) as a genotype by early-life nutrition interaction (GENI) ^11^. *Drosophila* provides exceptionally powerful tools and approaches for exploring the mechanisms underlying GENI at the single gene and genome-wide level ^12^. The *D. melanogaster* Genetic Reference Panel (DGRP) ^13^ consists of sequenced inbred lines derived from a natural population that has been extensively used to chart the genotype-phenotype architecture of complex traits, including behaviors and brain morphology ^12^. DGRP lines reared under different nutritional conditions show changes in behaviors and metabolic and transcriptional profiles, revealing a key role of GENI ^14–18^.

Alterations in sleep behavior are a common symptom of many neurological and psychiatric disorders, including neurodegenerative dementias and schizophrenia ^19, 20^. Therefore, uncovering the genes and pathways underlying the contribution of GENI to sleep behavior variation may shed light on altered neurodevelopmental mechanisms that can lead to mental illness. Few studies have focused on the role of early-life nutrition in sleep behavior. Prenatally malnourished rats exhibit decreased sleep and increased waking activity ^21, 22^; however, whether genetic variation affects such responses remains elusive. The influence of genetic variation on the effects of restrictive early-life nutrition on adult sleep behavior and brain structure remains unknown.

A genome-wide association (GWA) study of sleep using the DGRP ^23^, allowed the identification of naturally occurring sleep behavior-related genetic variants. Many of these variants were within or near candidate genes with human orthologs that have been associated with sleep, which suggests that genes affecting variation in sleep are conserved across species ^23^. Several of the candidate genes associated with natural variation in sleep affect developmental processes and neural function ^23, 24^ supporting the idea that variation in sleep is influenced by variation in the brain structures that control it.

The *Drosophila* mushroom bodies (MBs), an associative memory center, play an important role in sleep regulation ^25–27^. Early work showed that chemical ablation of the MBs results in a significant decrease in sleep ^25, 26^. MBs contain two types of sleep-regulating neurons: those that promote sleep when cyclic-AMP-dependent protein kinase A (PKA) is increased and those that inhibit sleep under such conditions ^26^. Each MB consists of around 2000 Kenyon cells (KCs) whose axons are arranged in parallel arrays projecting into different lobular structures, the α and β, the α’ and β’ and the γ lobes ^28^. KCs are sequentially generated from four neuroblasts in each hemisphere ^29^ that start to proliferate in embryos at stage 13 and continue uninterrupted until adult eclosion ^29, 30^. Therefore, nutritional restriction during early-life stages is likely to impact the development of MBs ^31^. Interestingly, natural genetic variation in the length and width of the α and β lobes has been correlated with variation in aggression and sleep behaviors ^32^.

Here, we assessed the effect of GENI on adult sleep behavior and MB morphology in the DGRP and found significant effects on both. We used GWA analyses to identify genetic variants and top candidate genes underlying GENI in the variation of sleep traits. Many proteins encoded by candidate genes are expressed in the MBs and form conserved protein-protein interaction networks required for neural development, translation, endocytosis regulation, ubiquitination, and lipid metabolism. Since genetic variants may affect the function or expression of candidate genes, we asked whether decreased expression of candidate genes in the MBs or in all neurons modifies sleep and MB morphology in response to early-life nutritional restriction. We found that diminished expression of a group of genes required for neural development, transmission of the nerve impulse and splicing, in specific neuronal populations, modifies the response of sleep behavior and MB morphology to early-life nutritional restriction. Together, our results suggest that tissue-specific variation in the expression of genes controlling these processes underlie GENI in sleep behavior and MB morphology.

## Results

To determine the consequences of nutritional restriction during early life on sleep behavior and brain morphology variation, we first evaluated whether reducing larval nutrients to 20% (restricted food, RF) of the standard culture medium (normal food, NF) affects development by analyzing adult size (S1 Fig). Flies reared in RF show a reduction in the notum length (NL) (S1C and S1F Figs), while the interocular distance (IOD) is not affected (S1B and S1E Figs) ^33^. These data confirm that nutrient restriction during development reduces the size of the thorax but not the head ^31^, which is evident in the increased IOD/NL ratio in flies reared in RF (S1D and S1G Figs).

We raised larvae of 73 DGRP lines on NF or RF and transferred newly eclosed adults to NF for three days to evaluate sleep behavior. We quantified nine sleep traits: total sleep duration, day and night sleep duration, day and night sleep bout number, day and night average sleep bout length, waking activity, and latency (i.e., the time it takes for the flies to start their first sleep bout after the lights are off) (Fig 1; S2 and S3 Figs; S1 and S2 Tables). We observed significant variation in sleep traits among the DGRP lines reared under both nutritional conditions (Fig 1; S2 and S3 Figs; Table 1; S1-S3 Tables).

**Fig 1.**
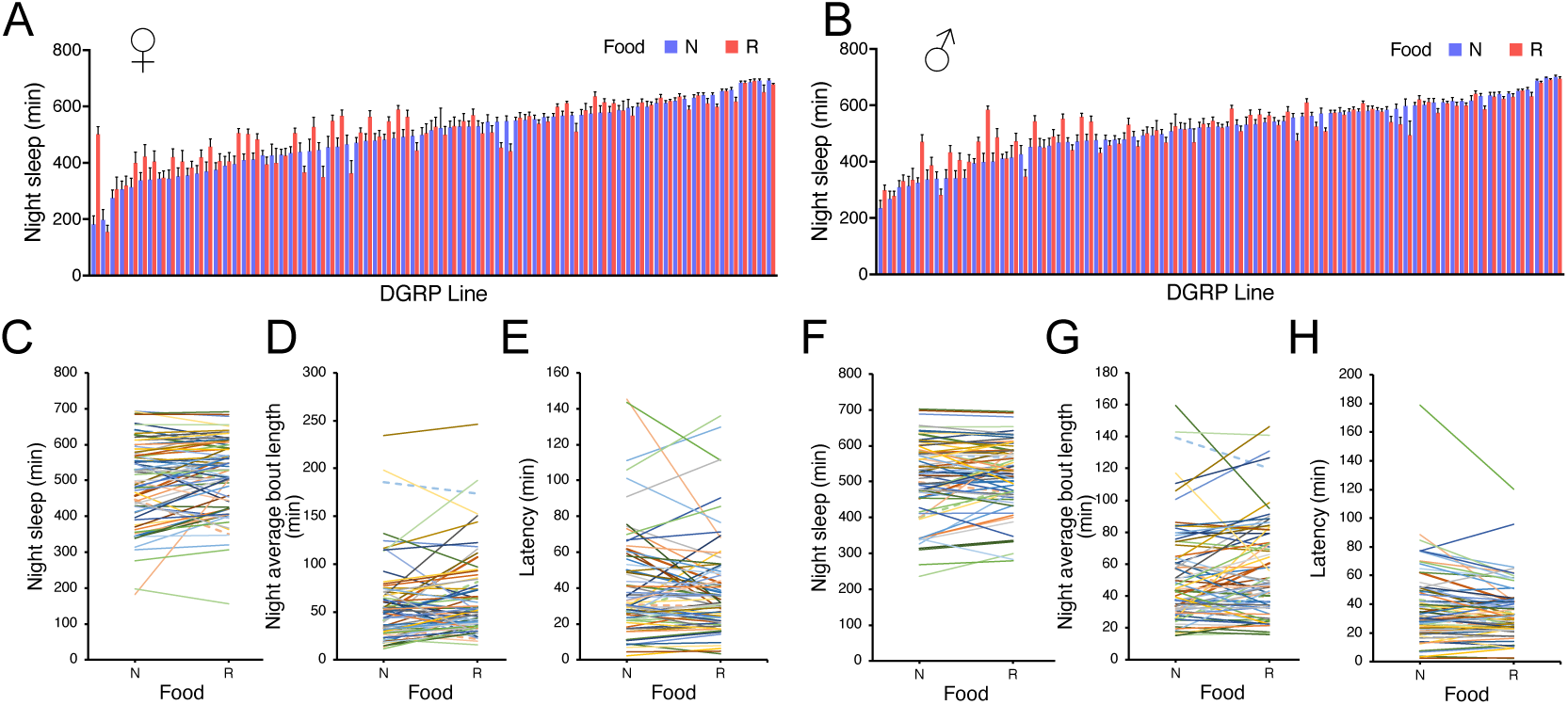
Sleep trait response to early-life nutrition. Histograms of sleep traits mean + SEM for night sleep trait in (A) females and (B) males. Reaction norms for sleep traits: (C) night sleep, (D) night average bout length and (E) latency in females; (F) night sleep, (G) night average bout length and (H) latency in males. Each DGRP line is represented by a different color. N: Normal food; R: Restricted food.

**Table 1.**
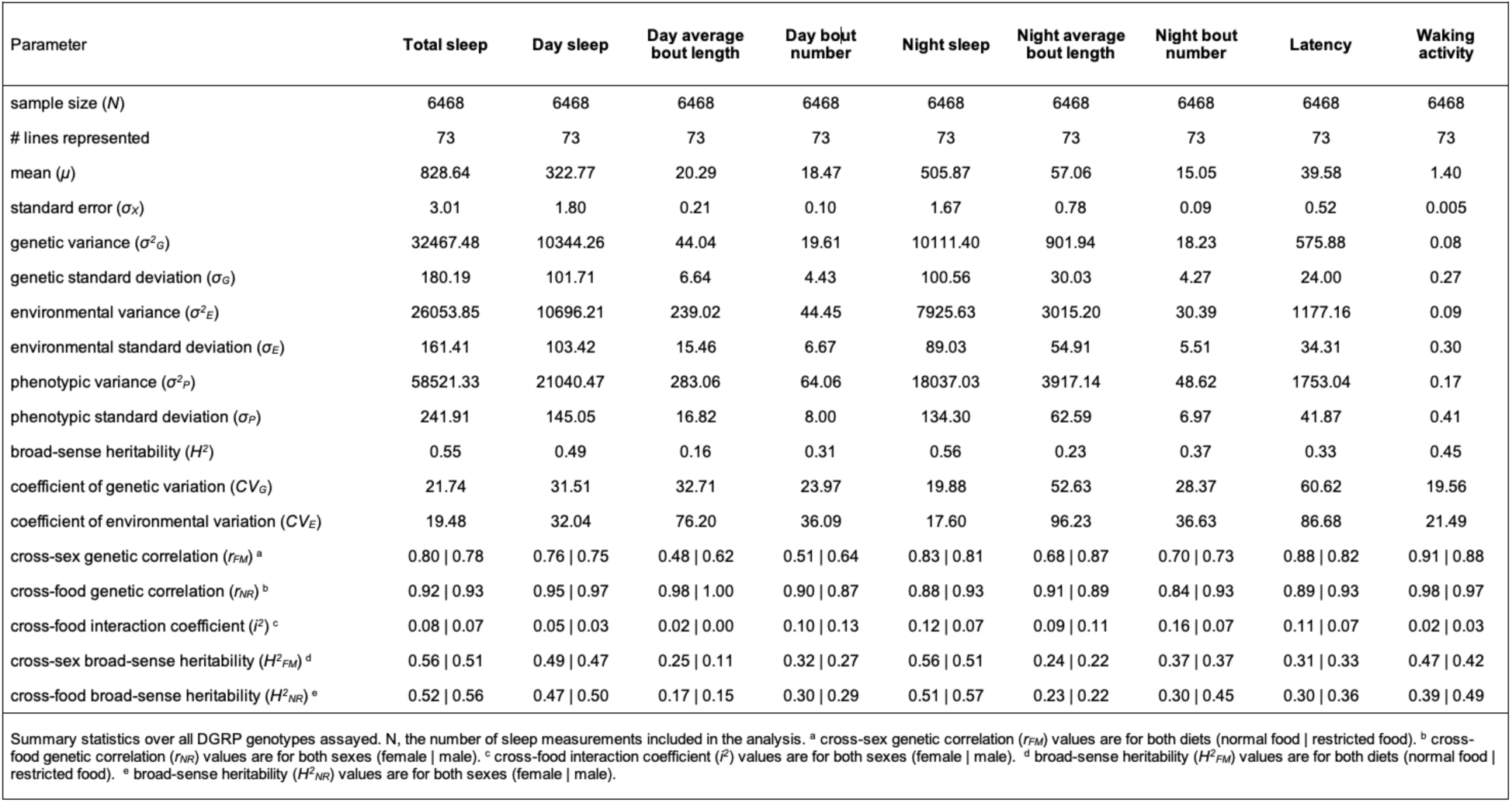
Quantitative genetic analysis of sleep traits on early-llfe nutrition.

Cross-sex genetic correlations (*r_FM_*), which represent the extent to which the same variants affect a trait in females and males, were significantly different from unity (NF, *r_FM_* = 0.48-0.91; RF, *r_FM_* = 0.62-0.88) (Table 1 and S3 Table). Thus, some polymorphisms affect sleep susceptibility to rearing diet in both sexes, while others will have sex-specific or sex-biased effects.

The differential responses of different genotypes to RF are evident from the complex pattern of crossing reaction norms, which is a hallmark of genotype by environment interaction (Fig 1; S2 and S3 Figs). To quantitate the contribution of GENI to the genetic variance, we estimated the interaction coefficient (*i*^2^) across food, which is calculated by subtracting the cross-environment genetic correlations from 1 (1- *r_NR_*). Estimates for *i*^2^ showed GENI contribution to night sleep traits genetic variation range from 9% in night bout length to 16% in night bout number in females, and from 7% in night sleep to 11% in night bout length in males. In contrast in day sleep traits, GENI contribution ranges from only 2% in day bout length to 10% in day bout number in females, and from 3% in day sleep to 13% in day bout number in males with no contribution to day bout length (Table 1 and S3 Table). Therefore, we decided to exclude day bout length from further analyses.

These data support a role of GENI in sleep variation, which may depend on variation in genes that act during development or whose adult expression is programmed during development in response to nutrient restriction.

### GENI contributes to morphological variation of the mushroom bodies

To assess to what extent GENI contributes to variation of MB morphology, we raised 40 DGRP lines under both dietary conditions during larval stages and examined the MB morphology of adult females. These include 36 lines that were previously used to demonstrate the natural variation of MB morphology ^32^. First, we assessed the gross morphology of α and β lobes ^32^. We observed a variety of large morphological defects at a broad range of frequencies (5–80%) (Fig 2; S4 Table), including missing or very thin structures and lobe fusions. These gross defects have been attributed to mutations that have major effects on MBs morphology ^32^, or could be due to multiple variants in each line. Strikingly, a total of 14 (35%) and 6 (15%) DGRP lines exhibited a decrease of the α- and β- lobe defects, respectively, when reared on restricted food (Fig 2; S4 Table). In turn, a total of 9 (22%) and 17 (42%) DGRP lines showed an increase of the α- and β- lobe defects, respectively, when reared on restricted food (Fig 2; S4 Table). These data indicate that early-life nutrition affects gross MB morphology.

**Fig 2.**
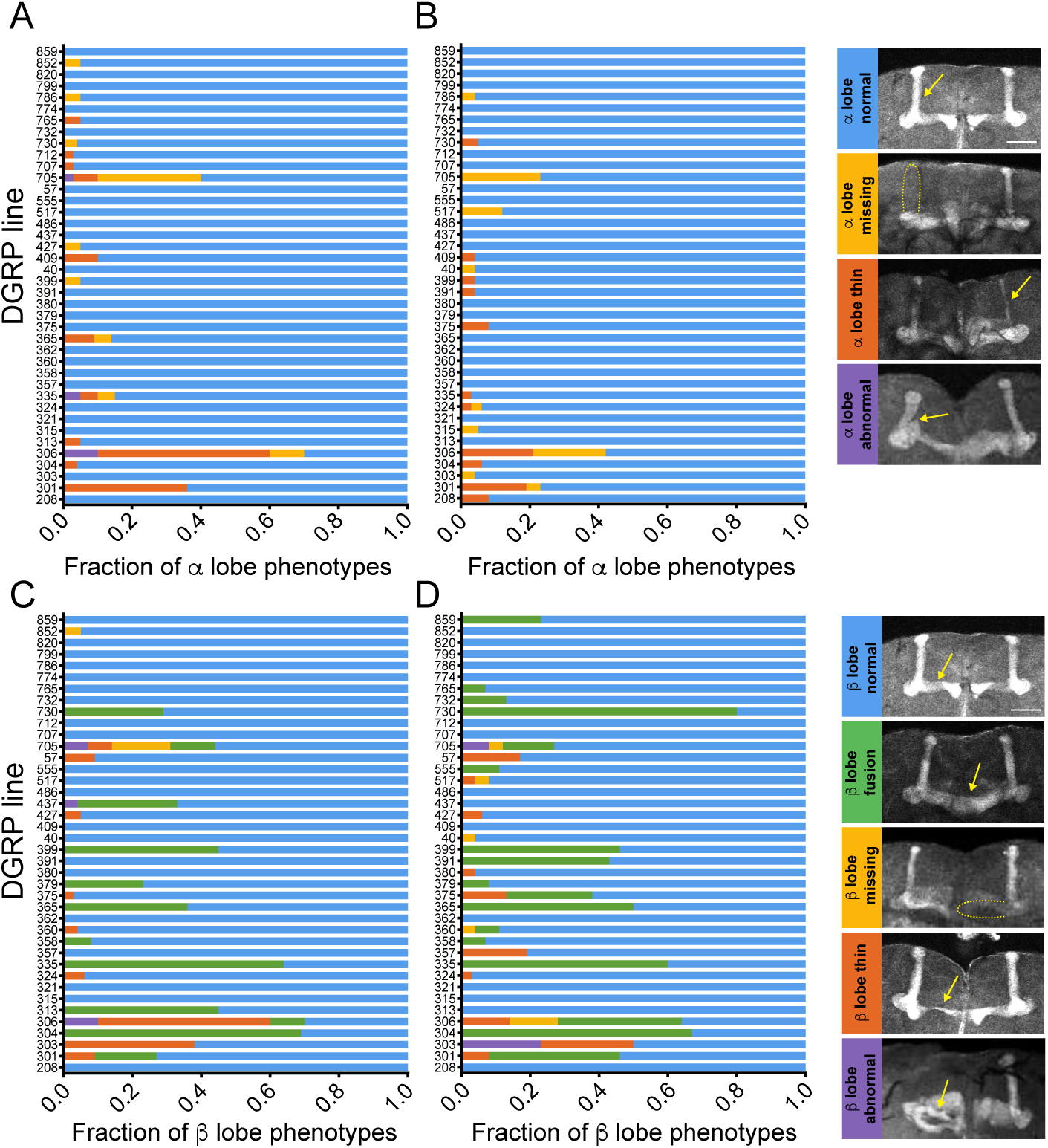
Gross morphological defects of MBs in the DGRP lines under prenatal nutritional restriction. Quantification of variation in gross MB defects of the 40 DGRP lines reared under (A, C) Normal or (B, D) Restricted food. (A, B) α-lobe phenotypes, (C, D) β-lobe phenotypes. Anti-Fas2 staining highlights the α- and β-lobes of the MBs in the adult brain of 3–5 days old females (scale bar, 50 µm). Categories of each MB phenotype are shown on the right side.

We next evaluated quantitative variation in MB morphology by measuring the length and width of the α and β lobes (Fig 3A; S5 and S6 Tables) ^32^ to reveal more subtle effects on the morphology of brain structures (Fig 3). Quantitative genetic analyses revealed substantial and significant genetic variation among the lines for length and width means of α and β lobes (Table 2; S7 Table). The contribution of genetic variation to phenotypic variation ranged from low for β-lobe length (*H*^2^ = 0.07 in NF and *H*^2^ = 0.13 in RF) to moderate for β-lobe width (*H*^2^ = 0.28 in NF and *H*^2^ = 0.30 in RF), α-lobe length (*H*^2^ = 0.23 in NF and RF) and α-lobe width (*H*^2^ = 0.31 in NF and *H*^2^ = 0.32 in RF) (Table 2; S7 Table).

**Fig 3.**
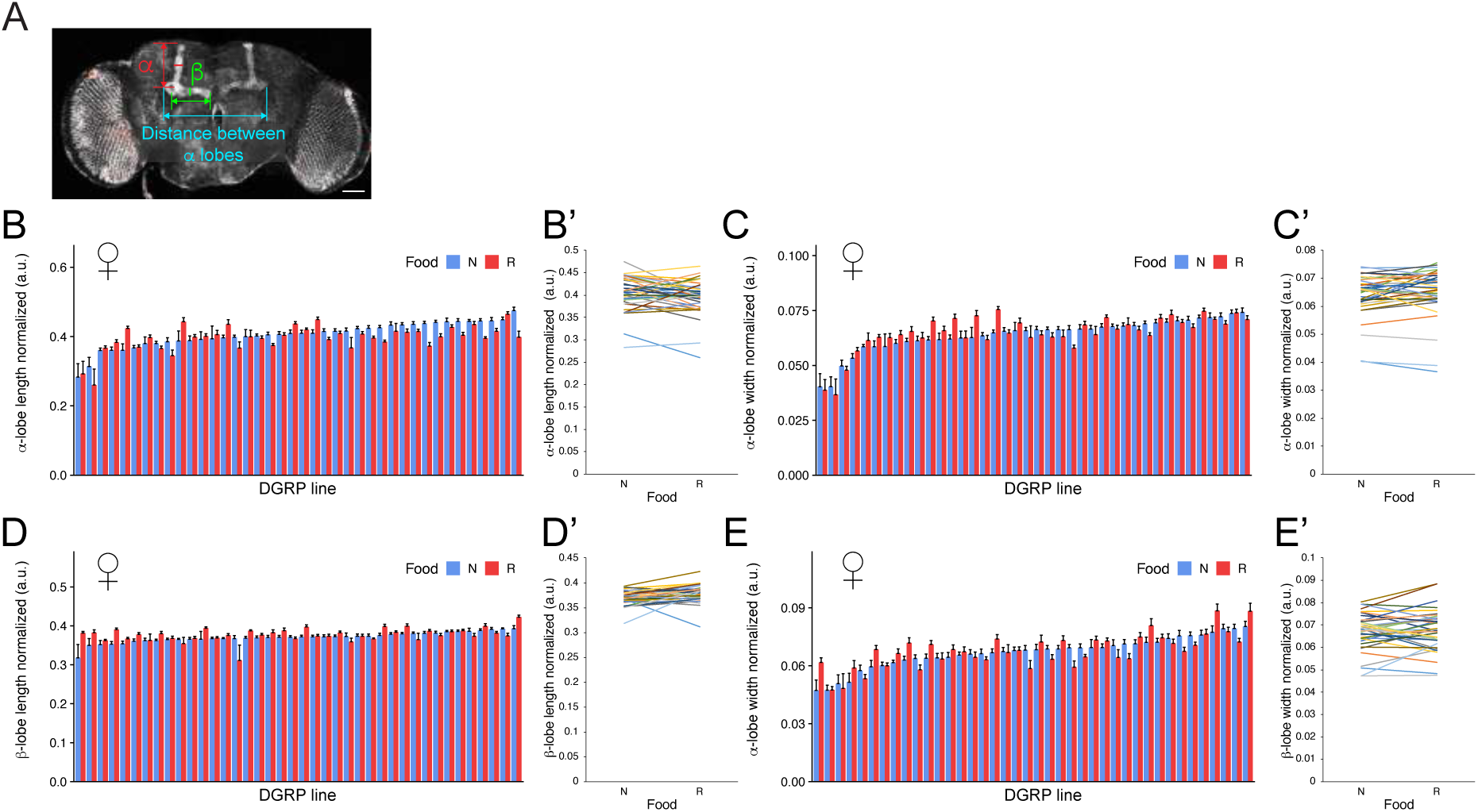
MBs morphometric variation in response to early-life nutrition. (A) Scheme showing morphometric measurements (scale bar, 50 µm). Morphometric measurements of normalized (B-B’) α-lobe length, (C-C’) α-lobe width, (D-D’) β-lobe length, (E-E’) β-lobe width. Histograms of 40 DGRP female lines reared under Normal (blue bars) or Restricted (red bars) food (B, C, D and E). Reaction norms for (B’) α-lobe length, (C’) α-lobe width, (D’) β-lobe length, (E’) β-lobe width. Each DGRP line is represented by a different color. N: Normal food; R: Restricted food.

**Table 2.** Quantitative genetic analysis of MB morphology traits on early-life nutrition.

| Parameter | α-lobe length | α-lobe width | β-lobe length | β-lobe width |
| --- | --- | --- | --- | --- |
| sample size ($N$) | 1743 | 1743 | 1743 | 1743 |
| # lines represented | 40 | 40 | 40 | 40 |
| mean ($\mu$) | 0.40 | 0.06 | 0.37 | 0.07 |
| standard error ($\sigma_X$) | 1.74E-03 | 3.23E-04 | 9.83E-04 | 3.58E-04 |
| genetic variance ($\sigma^2_G$) | 1.22E-03 | 5.77E-05 | 1.64E-04 | 6.50E-05 |
| genetic standard deviation ($\sigma_G$) | 0.03 | 0.01 | 0.01 | 0.01 |
| environmental variance ($\sigma^2_E$) | 4.03E-03 | 1.24E-04 | 1.52E-03 | 1.58E-04 |
| environmental standard deviation ($\sigma_E$) | 0.06 | 0.01 | 0.04 | 0.01 |
| phenotypic variance ($\sigma^2_P$) | 5.25E-03 | 1.82E-04 | 1.68E-03 | 2.23E-04 |
| phenotypic standard deviation ($\sigma_P$) | 0.07 | 0.01 | 0.04 | 0.01 |
| broad-sense heritability ($H^2$) | 0.23 | 0.32 | 0.10 | 0.29 |
| coefficient of genetic variation ($CV_G$) | 8.70 | 11.73 | 3.42 | 11.92 |
| coefficient of environmental variation ($CV_E$) | 15.81 | 17.20 | 10.40 | 18.59 |
| cross-food genetic correlation ($r_{NR}$) | 0.73 | 0.94 | 0.35 | 0.82 |
| cross-food interaction coefficient ($i^2$) | 0.27 | 0.06 | 0.65 | 0.18 |
| Summary statistics over all DGRP genotypes assayed. N, the number of morphology measurements included in the analysis. |  |  |  |  |

We found a significant line by food interaction term (*L x F*) for all four traits, indicating that flies with different genotypes respond differently to RF (Fig 3A’-D’, S7 Table). The contribution of GENI to morphological traits, measured as *i*^2^, was low for α- lobe width (*i*^2^ = 0.06), moderate for α-lobe length (*i*^2^ = 0.27) and β-lobe width (*i*^2^ = 0.18), and high for β-lobe length (*i*^2^ = 0.65) (Table 2 and S7 Table), indicating that the contribution of GENI to genetic variation highly depends on the trait.

In summary, we found that GENI contributes considerably to variation of MB morphology, suggesting an essential role in development and structure of this brain region.

### Genetic variants associated with GENI for sleep

To identify genetic variants underlying GENI in sleep behavior and MB morphology, we performed GWA analyses for the difference of phenotypic values for each trait between the two diets. We also excluded traits which Q-Q plots did not show larger than expected numbers of *P*-values below 10^-5^. These include day and night bout number in females, day sleep in males, and total sleep and waking activity in both sexes, and all morphology traits (S4-S6 Figs).

All variants associated with sleep traits from both sexes were pooled together for subsequent analysis as variants associated with sleep susceptibility to early-life nutrition. We found a total of 1,162 variants across all sleep traits (at a nominal reporting threshold of *P* ≤ 10^-5^) that mapped in or near to 611 candidate genes (S7 and S8 Figs; S8 Table). Among these, 11% of such variants were located within coding sequences, while 47% were in introns and the 5’ and 3’ untranslated region (UTR), and the remaining 42% were classified as intergenic (more than 1 kb away from the gene body) (S9 Fig; S8 Table). 95 (16%) candidate genes are highly enriched for GO terms (*FDR* < 0.05) associated with the function and development of the nervous system, including nervous system development, neurogenesis, neuron differentiation and development, and axonogenesis (S10 Fig; S8 Table). Twenty percent (122 out of 611) of the candidate genes are expressed in MBs either during larval stages or in adults (S9 Table) ^34–37^, 9 have orthologs associated with aspects of human brain size, and 41 genes have orthologs associated with human sleep traits (S11 Fig; S8 Table). These results suggest that the effect of early-life nutrition on sleep behavior in adulthood occurs in part through the regulation of neurodevelopmental mechanisms.

To identify potential cellular processes and molecular pathways underlying GENI, we generated protein-protein interaction networks with proteins encoded by candidate genes using the STRING database, which considers interactions based on direct (physical) and indirect (functional) associations ^38^. These proteins connect through processes that include translation (e.g., RpS27A and RpL23), vesicular trafficking (e.g., drongo), ubiquitination (e.g., Trim9 and Kel), lipid metabolism (e.g., LpR1 and LpR2), neural development (e.g., Fas2), and protease activity (e.g., Fur1 and Mmp2) (Fig 4). We found a significantly enriched network (PPI enrichment *P*-value = 1.06 x 10^-5^) using a high confidence score (score ≥ 0.700) (Fig 4).

**Fig 4.**
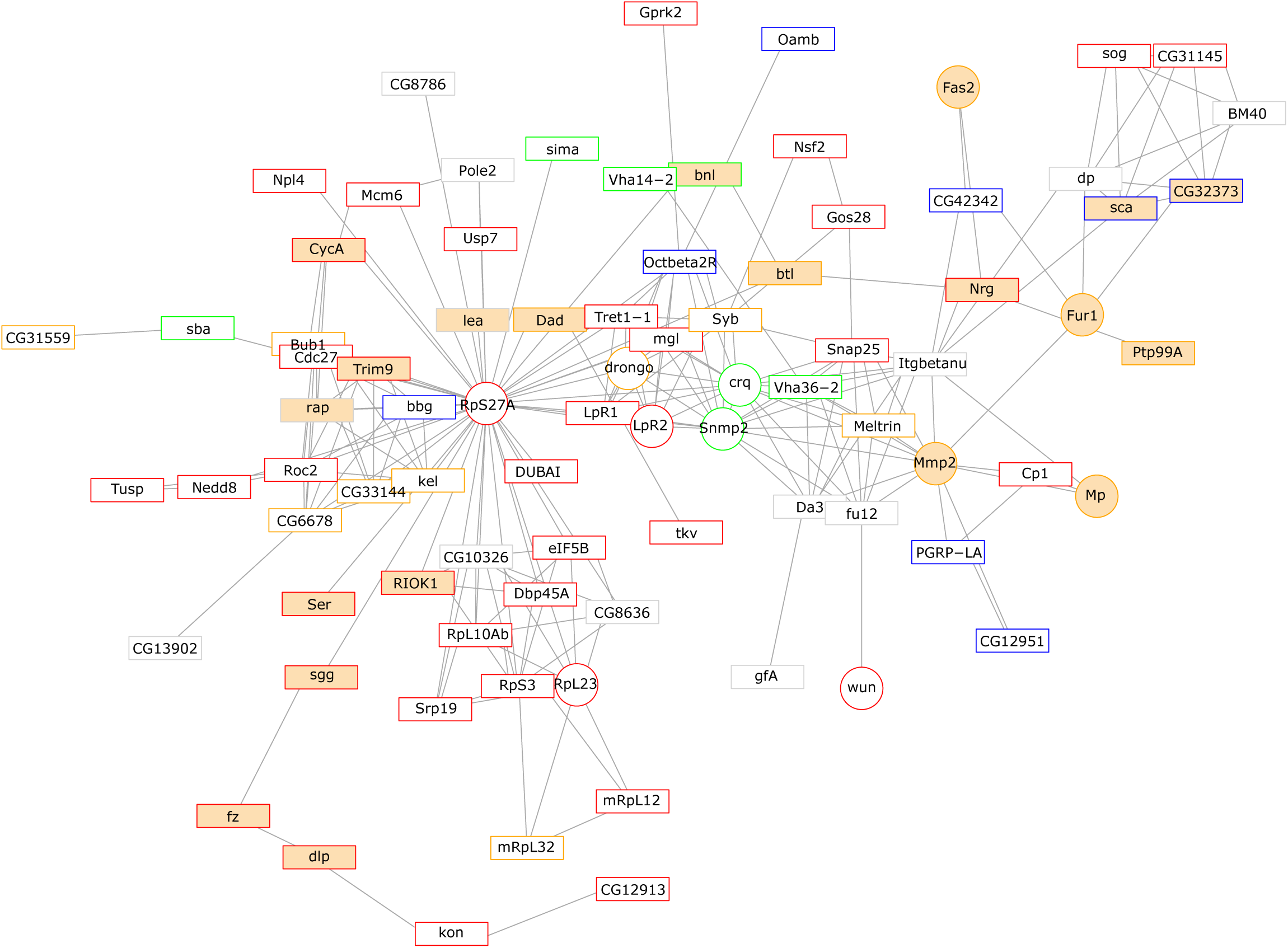
Protein-protein interaction network of proteins encoded by candidate genes underlying GENI in sleep behavior. Borders indicate the strength of the evidence for a human ortholog. Black: DIOPT score < 3; Blue: DIOPT score 3–6; Green: DIOPT score 7–9; Orange: DIOPT score 10–12; Red: DIOPT score 13–15. See S8 Table for the complete list of human orthologs and their DIOPT scores. Orange background indicates Gene Ontology enrichment category for Nervous System Development (GO:0007399). Circles denote genes that were validated by RNAi knockdown experiments.

### Functional assessment of candidate genes associated with GENI for sleep

We asked whether early-life nutrition modifies the knockdown effects of 17 candidate genes. These were chosen based on their role in nervous system development, our network analysis, or both (Figs 4 and 5; S10, S12-S15 Figs; S10 Table).

**Fig 5.**
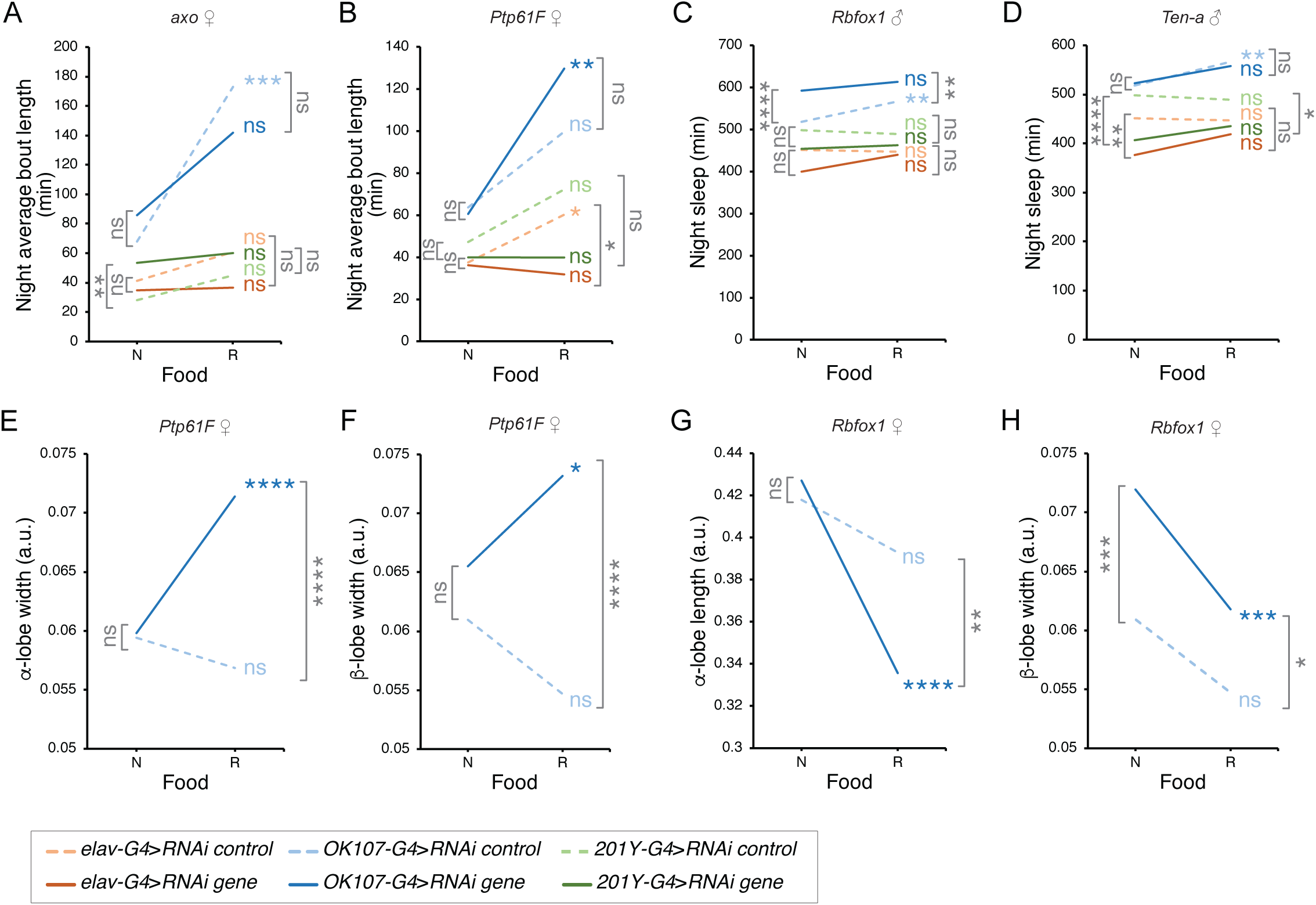
RNAi-mediated knockdown of candidate genes associated with GENI in sleep affects sleep traits and MBs morphology in response to early-life nutrition. (A-D) Sleep analyses of flies expressing candidate gene RNAis in the MBs or in all neurons in NF and RF. Reaction norms depict sleep trait mean in the two experimental conditions: RNAi control (dashed lines) and RNAi candidate gene (solid lines) using a pan-neural driver (*elav-Gal4*, orange lines) or MBs driver (*OK107-Gal4*, blue lines; *201Y-Gal4*, green lines) from flies reared under Normal or Restricted food. (E-H) MBs morphology analyses in response to early-life nutrition after knockdown of candidate genes in the MBs. Reaction norms showing means of female MB morphology traits in the two experimental conditions: RNAi control (dashed lines) and RNAi candidate gene (solid lines) using a MB driver *OK107-Gal4* from flies reared under Normal or Restricted food. We performed two-way ANOVA followed by Bonferroni’s post hoc test (*p≤ 0.05; **p≤0.01; ***p≤0.001; ****p<0.0001; ns non-significant) to compare the effect of control or candidate gene RNAi knockdown between normal and restricted food (see color coded asterisks and ns). We also performed (A) Welch’s t-test or (B-H) one-way ANOVA followed by Dunnett’s post hoc test (*p≤ 0.05; **p≤0.01; ***p≤0.001; ****p<0.0001; ns non-significant) to compare the effect of control versus candidate gene RNAi knockdown in each food condition (see gray coded asterisks and ns). The candidate gene name and sex of adult flies are indicated at the top of each plot. N: Normal food; R: Restricted food.

To test the hypothesis that these genes affect GENI in MBs we reduced their expression from embryonic stages onwards in all MB neuroblasts by driving the expression of specific *UAS*-*RNAi* transgenes using the *OK107-Gal4* driver (Fig 5; S12- S13 Figs; S10 Table) ^35^. We use the *201Y-Gal4* driver for genes that showed an effect with *OK107-Gal4*, as the latter is strongly expressed in other brain regions, including the optic lobe, pars intercerebelaris (PI), tritocerebrum (TR), and subesophageal ganglion (SOG) (Fig 5; S15 Fig; S10 Table). *201Y-Gal4* displays a more restricted pattern both in the MB, specifically in the γ and α/βc and neurons, and to a lesser extent in the PI, TR, and SOG ^39^.

In addition, to control for the specificity of the RNAi-mediated knockdown, we use a second RNAi for candidates that showed effects with *OK107-Gal4* (S14 Fig; S10 Table). To distinguish whether the effects on sleep are specific to MBs, other regions of the nervous system, or both, we also drove the expression of *UAS-RNAi* transgenes in all neurons using the *elav-Gal4* driver (Fig 5; S12-S15 Figs; S10 Table) ^40^. We reared flies expressing the RNAi and their specific control genotypes in NF or RF and quantified sleep behavior in adults (Fig 5; S12-S15 Figs; S10 Table).

All candidate gene RNAis tested affected at least one sleep trait in response to early-life nutrition in males, females, or both sexes, suggesting pleiotropy (S13 Fig). Interestingly, early-life nutrition modifies the effects of the expression of the RNAis with *elav-Gal4* in 33 traits, while with *OK107-Gal4* in 57 traits, supporting a key role of the MB in GENI (S13 Fig). The knockdown of 7 genes affects the same trait associated with the candidate genetic variant, supporting their role as specific modifiers of behavioral response to diet (Fig 5, S13 Fig). Among them *Tenascin-a* (*Ten-a*) ^41^, *axotactin* (*axo*) ^42^, *guk-holder* (*gukh*) ^43^, *Multiplexin* (*Mp*) ^44^, *Protein tyrosine phosphatase 61F* (*Ptp61F*) ^45^, and *wunen* (*wun*) ^46^, affected more traits than in the GWA analyses. On the other hand, early-life nutrition only modifies *RNA-binding Fox splicing factor 1/ Ataxin 2-binding protein 1* (*Rbfox1)* ^47^ effect in night sleep, supporting their specific role in this trait.

In females, the expression of an *RNAi* targeting *axo* (*axo-RNAi*) with *OK107- Gal4*, a gene required for transmission of the nerve impulse ^42^, suppresses night average bout length increase in response to early life nutrition (Fig 5, S13 Fig, S10 Table). We found a similar response when *axo-RNAi* was driven with the *201Y-Gal4* driver, supporting its specific role in the MB. Here, RF suppresses the difference in phenotypic values between the control and the RNAi expressing flies observed in NF (Fig 5A). The expression of a second axo-RNAi (*axo (2)-RNAi*) showed similar but not significant effects (S14A and S15 Figs). These data suggest that variation in *axo* expression in the MB may modify the adult’s response to early-life nutrition restriction.

Unlike *axo-RNAi*, the expression of *Ptp61F-RNAi* in the MBs with *OK107-Gal4* increases night average bout length in response to RF, and suppresses it when expressed with *elav-Gal4*, suggesting that its expression in different neuronal populations may affect in different ways sleep response to early-life nutrition restriction. Although *Ptp61F* second RNAi (*Ptp61F (2)-RNAi*) did not show similar effects in response to diet (compare Fig 5B with S14B and S15 Figs) we decided to continue our work using *Ptp61F-RNAi* since its functionality has been proven in hemocytes, where it promotes activation of Jak/Stat ^48^.

Analysis of night sleep shows that expression of *Rbfox1-RNAi* or *Ten-a*-*RNAi* under the control *OK107-Gal4* slightly suppresses the night sleep increase in RF in males. In addition, expression of *Ten-a RNAi* under the control of *elav-Gal4*, decreases night sleep in NF but not in RF (Fig 5, S13 Fig, S10 Table). Expression of a second *Rbfox1-RNAi* (*Rbfox1 (2)-RNAi*) under the control of *OK107-Gal4* phenocopies the *Rbfox1-RNAi* effect (S14C and S15 Figs, S10 Table). These data support the hypothesis that variation of the expression of *Rbfox1* in cells of the MBs that only expresses *OK107-Gal4* but not *201y-Gal4* underlies GENI in night sleep behavior. On the other hand, although *Ten-a (2)-RNAi* did not phenocopy the effect of *Ten-a RNAi*, we continue using the latter since the pan-neuronal expression of *Ten-a-RNAi* results in undetectable levels of Ten-a immunostaining, and that the function of *Ten-a (2)-RNAi* has not been tested ^49^ (S14D and S15 Figs, S10 Table).

Interestingly, pan-neuronal expression of and RNAi targeting *gukh,* a gene involved in cell polarity and synapse formation ^43^, increases night sleep in response to early-life nutrition restriction (S12A and S13 Figs, S10 Table). In contrast, its expression under the control of *OK107-Gal4* increases sleep in both diets and suppresses the increase in sleep in response to RF observed in the control genotype. These data suggest that *gukh* expression in different neuronal populations affects the response to early-life nutrition restriction in different ways, similar to *Ptp61F-RNAi* (S12A and S13 Figs, S10 Table). On the contrary, knockdown of *Mp* ^44^ using *elav-Gal4* suppresses the increase in night sleep observed in the control genotype (S12B and S13 Figs, S10 Table).

Finally, expression of an RNAi targeting the lipid phosphatidate phosphatase *wunen* (*wun*) under the control of *OK107-Gal4* in females increases latency in response to RF, while in males, it increases latency in both diets (S12C, S12D and S13 Figs, S10 Table). Moreover, its knockdown using *elav-Gal4* in females increases latency to the same extent in NF and RF, while in males, it decreases it in response to RF (S12C, S12D and S13 Figs, S10 Table). These data indicate that the interaction between *wun* knockdown and early-life nutrition depends on the sex and neuronal population affected and suggest that early-life nutrition modifies the effect on sleep of diminished expression of *wun* in non-MB neurons.

In summary, we show that reducing candidate gene expression in all neurons or specifically in MBs, can have different effects on sleep behavior upon early-life nutrition and that these effects can be sex-specific. Moreover, depending on their local expression in the nervous system, candidate genes may be involved in different aspects of sleep regulation in response to early-life nutrition. These data support the idea that a subgroup of candidate genes associated with variation in sleep behavior act at the MBs underlying GENI in sleep behavior.

### Early-life nutrition restriction modifies the effect of *PTP61F* and *Rbfox1* knockdown in MB morphology

Since early-life nutrition modifies the sleep effects associated with *axo*, *Ptp61F, Rbfox1*, and *Ten-a* knockdown in the MB, we asked whether they affect MB morphology and whether their effects are modified by early-life nutrition. We used the *OK107-Gal4* driver since it is strongly expressed in all MB neurons ^39^.

We found that α-lobe width and β-lobe width increases in response to RF in flies expressing *Ptp61F-RNAi* under the control of *OK107-Gal4*, but not in control flies (Fig 5A and 5B, S16 Fig, S11 Table). On the other hand, the expression *Rbfox1-RNAi* decreases α-lobe length in response to RF, while the control genotype does not display a response to diet, resulting in a significant length reduction RF (Fig 5C, S16 Fig, S11 Table). In addition, *Rbfox1 knockdown* increases the α-lobe width in RF and β-lobe length in NF, however, these genotypes as the control did not show differences between diets (S16 Fig, S11 table). Finally, *Rbfox1* knockdown increases β-lobe width in NF and RF (Fig 5H, S11 table), but both the control and *Rbfox1-RNAi* expressing flies decrease β-lobe width in response to RF, indicating plasticity but not GENI (Fig 5, S11 table). Together, these data indicate that variation in the expression of *Ptp61F* or *Rbfox1* in the MB neurons interacts with early-life nutrition affecting MB morphology.

Finally, we found that the expression of *Ten-a-RNAi* and *axo-RNAi* did not show clear GENI effects. Their knockdown increases α-lobe width in both diets (S16 Fig, S11 table) and *axo* knockdown also increases α-lobe width in response to RF indicating plasticity (S16 Fig, S11 table). *Ten-a* knockdown increases β-lobe length in RF but not in NF suggesting weak GENI. Finally, we did not find differences between the *axo* knockdown and the control in the other traits in response to diet (S16 Fig, S11 table).

## Discussion

The development and function of the central nervous system in animals, including humans, rely on proper nutrition during the prenatal period ^22, 50^. In humans, early-life nutritional restriction increases the risk of mental and neurological diseases associated with morphological brain defects ^51, 52^. To what extent the interaction between the genotypes of individuals interacts with early-life nutrition to sensitize or protect from developing pathological behaviors is unknown. The use of animal models and genetic reference panels in mammals and *Drosophila*, such as the DGRP, have been critical to determining the contribution of GENI in complex behaviors at the genome-wide level^16, 17^.

Adult flies display genetic variation in the sensitivity of sleep behavior and MBs morphology to early-life nutrition. Moreover, we found that gross morphological defects displayed by a group of DGRP lines can be strongly modified by nutrition, indicating that early-life diet plays an important role in modifying the phenotype associated with single or multiple mutations that strongly affect morphology.

We identified candidate genes and molecular processes whose variation may support sleep behavior variation in response to the environment. A total of 135 of the 611 candidate genes (22%) associated with GENI for sleep phenotypes were found in a previous analysis of sleep variation in the DGRP under standard nutritional conditions ^23^ (S16 Table), and only 12 out of the 611 have been previously linked to sleep phenotypes ^53^. Thus, the contribution of GENI to variation in sleep behavior involves a new set of genes.

Although the study of only 73 lines may lead to a higher rate of false positives ^54^ and to miss interesting candidate genes due to the low frequency of some variants, we were able to find a significant protein-protein interaction network that reveals biological processes that affect GENI ^38^. We identified a subset of cellular processes involved in proteostasis, lipid metabolism, and nervous system development and function, suggesting that perturbations in the proper expression of genes encoding these proteins may impact central nervous system function later on. Cytosolic and mitochondrial ribosomal proteins generate a subnetwork including Rps27A, a ribosomal protein fused to a single copy of ubiquitin ^55^, and RpL23 ^55, 56^. Mutants of ribosomal proteins result in the Minute phenotype, characterized by delayed development due to reduced protein biosynthesis ^57, 58^. This condition decreases *Drosophila* insulin-like peptide (Dilps) secretion systemically by affecting the insulin-producing cells (IPCs) in a cell-autonomous manner, leading to reduced body size and delayed larval development ^59^. Our results showed that variation in the expression of *Rps27A* and *RpL23* in the MB or all neurons modify sleep behaviors in response to early-life nutrition (S13 Fig and S10 Table), suggesting that modulation of protein biosynthesis can play a neuronal-specific role in GENI. Since neuronal knockdown *Rps27A* or *RpL23* does not affect latency or night sleep, to whose variation the original genetic variants are associated, we support the hypothesis that they play a role in non-neuronal cells, such as in the fat body, contributing to GENI more systemically.

Rps27A is a central network node connecting the ribosomal proteins subnetwork with other subnetworks containing the ubiquitin ligases and the lipid receptor proteins LpR1, LpR2, and Megalin ^60^. LpR1 is required in a group of octopaminergic neurons to enhance starvation-induced hyperactivity by inhibiting the degradation of the adipokinetic hormone receptor ^61^. Here we found that *LpR2-RNAi* expression with *OK107-Gal4* driver suppresses the increase in night sleep in response to early-life undernutrition in males and the decrease of night bout numbers in females (S13 Fig and S10 Table).

The network also contains a group of proteins involved in axon growth, targeting, synaptogenesis, and fasciculation, including PTP99a ^62^, Nrg ^63^, and Fas2 ^64, 65^. Another group is enriched in protein degradation, highlighting the ubiquitin ligase encoded by Trim9, which mediates Netrin function in axon guidance ^66^. These data suggest that variation in the expression of genes required for neuronal terminal differentiation contributes to GENI in sleep behavior.

Recent GWA studies on sleep traits and insomnia in humans provide insight into the genetic basis for variation in sleep ^67–70^. We found that 41 genes identified in these GWA studies, including *FURIN* and *RBFOX1*, are human orthologs of *Drosophila* genes associated with different sleep phenotypes in response to early-life nutrition (S14 Table). For example, Furin convertase is an enzyme that processes the precursor of endothelin-1 (ET-1) ^71^ and is associated with insomnia ^68^. Furin convertase also cleaves the precursor form of brain-derived neurotrophic factor (BDNF) to generate mature BDNF ^72^. BDNF plays a role in homeostatic sleep regulation ^73^ and an essential role in brain development and synaptic plasticity ^74^.

Early-life nutrition restriction modifies the knockdown effects of two regulators of Insulin signaling in sleep behavior and brain morphology: *Ptp61F* ^75^ and *Rbfox1* ^76^. *Ptp61F* encodes a protein tyrosine phosphatase ortholog to the mammalian PTP1B, which dephosphorylates insulin receptor (IR), inactivating it ^45^. Since overexpression of insulin/IGF-like peptides in glia or PI3K/Akt activation in neuroblasts results in increased proliferation under protein restriction ^77^, we propose that knockdown of *Ptp61F* contributes to uncoupling neuroblast growth from systemic control. These effects may increase the number of adult neurons and α- and β-lobe width.

On the other hand, *Rbfox1* encodes a conserved RNA-binding protein with nuclear isoforms that regulate tissue-specific alternative splicing ^76^, while cytoplasmic isoforms regulate mRNA translation ^78, 79^, suggesting that variation in these processes underlies sleep and neuronal morphology sensitivity to early-life nutrition. Rbfox1 targets alternative splicing of Tsc2 ^80^, which antagonizes cell growth and cell proliferation induced by Insulin signaling in *Drosophila* ^81^. On the other hand, Insulin signaling inhibits Tsc1 and Tsc2 to promote axon growth and branching during metamorphosis ^82^. Accordingly, our results suggest that variation in *Rbfox1* expression interacts with early-life nutrition to affect α-lobe length, which may be due to axon growth defects.

Cytoplasmic Rbfox1 regulates the expression of synaptic genes by binding to the 3’UTR of mRNAs that are targets of microRNAs, independent of its effect on splicing ^79^. Moreover, disruption of *Rbfox1* in the central nervous system leads to neuronal hyperactivity, while the deletion of *Rbfox2* results in cerebellum development defects in mice ^83, 84^. Thus, the mechanism by which variation in *Rbfox1* expression affects sleep may also be linked to its role in synapse formation.

DGRP lines reared under different nutritional conditions showed changes in the metabolic phenotype and transcriptional profiles that rely on genotype by environment interactions ^14–18, 85^. These metabolic changes correlate with previously reported phenotypes on the DGRP ^14^. Therefore, polymorphisms in *Ptp61F* and *Rbfox1* could significantly impact the response to environmental cues that affect nervous system development and function.

Our results suggest that gene expression variation in the MBs neurons and other groups of neurons can have different effects in sleep behavior and brain morphology depending on the nutritional environment during development. Our work supports the hypothesis that variants affecting local gene expression in the brain play an essential role in GENI in sleep behavior. It also may explain why few genetic variants associated with GENI in complex behaviors affect gene expression in the whole animal ^16^. Further evaluation of the effect of genetic variants in candidate gene expression ^86^ in specific developing and adult brain regions and neural lineages will be necessary to unveil the molecular and cellular mechanisms underlying GENI in sleep behavior.

## Materials and Methods

### *Drosophila* stocks and husbandry

We used 74 DGRP lines ^13^. The DGRP lines and *GAL4* drivers were obtained from the Bloomington *Drosophila* stock center (http://flystocks.bio.indiana.edu/). *UAS*-RNAi transgenic flies were obtained from Vienna *Drosophila* RNAi Center (VDRC) (https://stockcenter.vdrc.at), and the Transgenic RNAi Project (TRiP) at Harvard Medical School (http://www.flyrnai.org). All fly lines used are listed in S17 Table. All flies were reared under standard culture conditions (25 °C, 60–70% humidity,12-hour light:dark cycle) and controlled density.

### *Drosophila* culture media

The Normal Food (NF) diet for stock maintenance contains: 10% (w/v) Brewer’s yeast, 5% (w/v) sucrose, 1.2% (w/v) agar, 0.6% (v/v) propionic acid, 3% (v/v) nipagin ^87^. The Restricted Food (RF) medium contains 20% of Brewer’s yeast and sucrose of NF (2% (w/v) Brewer’s yeast, 1% (w/v) sucrose, 1.2% (w/v) agar, 0.6% (v/v) propionic acid, 3% (v/v) nipagin) ^87^. For each DGRP line, we used 35 females and 15 males to obtain comparable levels of offspring density. For the few DGRP lines that yield less offspring in restricted food than in normal food, we used 50 females and 20 males. Adult flies were discarded after three days of egg-laying on NF or RF. Newly eclosed adults were transferred to new NF vials for three days before any behavioral assay.

### *Drosophila* morphometric body measurements

We evaluated two morphometric body traits in adult Oregon-R flies reared under normal or restricted food. We distributed approx. 100 eggs in bottles of NF or RF in two biological replicates. Flies were reared under standard culture conditions (25 °C, 60– 70% humidity,12-hour light:dark cycle). After eclosion, flies were placed under NF for 3- 5 days. Adult females and males were selected and fixed for 24 h in 70% ethanol. The flies were placed on Sylgard plates with 70% ethanol, legs and wings were removed avoiding damage. After immobilizing the sample, the ILUMINA software was used to take a picture of the head and thoraxes at 6X with a stereoscopic magnifier attached to an INFINITY Lumenal Photo Camera 1 ^88^. All images were processed and assembled using Fiji software and Adobe Illustrator 2020. Depending on the number of flies available, between twenty and twenty-five flies per sex were measured. Interocular distance (IOD) was measured from eye edge to eye edge along the anterior edge of the posterior ocelli and parallel to the base of the head (S1A Fig) ^33^. Notum length (NL) was measured in the midline from the anterior edge of the thorax to the anterior edge of the scutellum (S1A Fig).

### Sleep phenotypes

We evaluated sleep traits in 73 DGRP lines (S1 Table). We picked groups of 3 to 10 DGRP lines to grow them in bottles in the two diet conditions without any block design plan. After eclosion, adult flies (males and virgin females) were transferred to vials with NF for 3 days until the sleep behavior was assessed. All sleep measurements were performed under standard culture conditions (25 °C, 60–70% humidity,12-hour light:dark cycle). Sleep measurements were replicated three times for most lines. Eight flies of each sex and each diet (NF and RF) were measured in one *Drosophila* Activity Monitors (DAM2, Trikinetics, Waltham, MA) per replicate. To mitigate the effects of both social exposure and mating on sleep behavior, males and virgin females were collected from each line and retained at 30 flies per same-sex same-diet vial. Individual flies were loaded into DAM2 monitors and sleep and activity parameters were recorded for seven continuous days. To mitigate the effects of CO_2_ anesthesia or any other potential acclimation effects, the first two days of data recording were discarded. The DAM2 monitors use an infra-red beam to detect activity counts in individual flies as they move past it; five minutes without an activity count is defined as sleep ^89, 90^. Flies were visually examined after the sleep and activity recordings were completed; data from any flies that did not survive the recording period was discarded. All DGRP sleep behavior analysis was done with data collected from days 3 to 4 after flies were placed into the DAMs. PySolo ^91^ software was used to calculate nine sleep parameters: total, night and day sleep duration in minutes, night and day sleep bout number, night and day average sleep bout length; it also calculated sleep latency, the time in minutes to the first sleep bout after lights are turned off, and waking index, the average number of beam crossings within an active bout.

### Immunohistochemistry

We used 40 DGRP lines based on lines previously tested for MB morphology (36 lines in common) ^32^, which were also used to evaluate sleep behavior (39 lines were contained in the 73 lines evaluated) (S17 Table). Adult brains from female flies reared on NF or RF under standard culture conditions (25 °C, 60–70% humidity,12-hour light:dark cycle) were dissected and processed for immunohistochemistry as described previously ^32, 92^. All flies were between 3 and 5 days old at the time of dissection.

*Drosophila* brains were fixed in phosphate-buffered saline (PBS)-4% formaldehyde for 25 min at room temperature, washed 3 times with PBS-0.3% Triton X (PBST), and blocked in PBST containing 5% Normal Donkey Serum for 30 min at room temperature. Brains were incubated overnight at 4°C with mouse monoclonal anti-Fasciclin 2 antibody (1D4) (1:10; Developmental Studies Hybridoma Bank, University of Iowa, IO, USA) to visualize mushroom body α and β lobes. After washing 3 times with PBST, brains were incubated with a 100-fold dilution of Rhodamine (TRITC) AffiniPure donkey anti-mouse IgG for 2 h at room temperature, followed by washing 3 times with PBST. Brain samples were mounted in Vectashield mounting medium (Vector Laboratories, Burlingame, CA). Immunostaining was documented with an Olympus Fluoview FV1000 confocal microscope. To avoid any effects of variation in Fas2 expression between DGRP lines, we adjusted fluorescence intensities so that unambiguous measurements could be made ^32, 92^.

### Morphometric measurements

The length and width of the MB α and β lobes in adult female brains were evaluated using the straight-line measurement tool from Fiji/Image J software (version 2.1.0/1.53c) ^93^ and expressed as values relative to the distance between the α lobe heels as described previously ^32, 92^. This internal calibration controls for differences in brain size when assessing variation in morphometric parameters among genotypes. MBs were scored individually (*i.e.*, per hemisphere). Values were obtained for 10-12 brains for all genotypes, thus allowing the analysis of 20-24 hemispheres. Images, diagrams, and figures were assembled using Adobe Photoshop 2020 and Illustrator 2020.

### Quantitative genetic analyses of sleep in the DGRP

We partitioned the variance in each sleep parameter in the DGRP using mixed model analyses of variance (ANOVA): *Y* = *µ* + *L* + *F* + *S* + (*L* × *F*) + (*L* × *S*) + (*F* × *S*) + (*L* × *F* × *S*) + *R*(*L)* + *F x R*(*L)* + *S x R*(*L)* + *F x S x R*(*L)* + *ε*, where *Y* is the sleep parameter; *µ* is the overall mean; *L* and *R* are the random effects of line and replicate, respectively; *F* and *S* are the fixed effects of food (control, restricted) and sex (males, females), respectively; and *ε* is the error variance. In addition, we performed reduced analyses within: (i) each food condition using mixed model ANOVAs of form *Y = µ + L + S* + (*L* × *S*) + *R*(*L)* + *S x R*(*L) + ε*; and (ii) each sex condition using mixed model ANOVAs of form *Y = µ + L + F* + (*L* × *F*) + *R*(*L)* + *F x R*(*L) + ε*. All ANOVAs were performed using the PROC GLM function in SAS.

We calculated broad-sense heritability by sex across food as *H*^2^ *=* (*σ*^2^*_L_ + σ*^2^*_LF_*)*/*(*σ*^2^*_L_ + σ*^2^*_LF_ + σ*^2^*_E_*), where *σ*^2^*_L_* is the variance component among lines, *σ*^2^*_LF_* is the line-by-food variance component, and *σ*^2^*_E_* is the residual variance. In addition, we calculated broad-sense heritability by food across sex as *H*^2^ *=* (*σ*^2^*_L_ + σ*^2^*_LS_*)/(*σ*^2^*_L_ + σ*^2^*_LS_ + σ*^2^*_E_)*, where *σ*^2^*_LS_* is the line-by-sex variance component. We calculated broad-sense heritability by food and sex as *H*^2^ *= (σ*^2^*_L_)*/(*σ*^2^*_L_ + σ*^2^*_E_*). We calculated genetic correlations by sex across food *r_NR_* = (*σ*^2^*_L_*)/(*σ*^2^*_L_ + σ*^2^*_LF_*), and by food across sex *r_MF_ =* (*σ*^2^*_L_*)/(*σ*^2^*_L_ + σ*^2^*_LS_*). We defined the interaction coefficient across food (*i*^2^) as a measurement of the genotype by early-life nutrition interaction (GENI) contribution to the genetic variation, *i*^2^ = 1- *r_NR_*.

### Quantitative genetic analyses of morphometric measurements in the DGRP

We partitioned the variation in the length and width of the α and β MB lobes in the DGRP using mixed model ANOVA: *Y* = *µ* + *L* + *F* + (*L* × *F*) + *ε*, where *Y* is the morphometric parameter; *µ* is the overall mean; *L* is the random effect of line; *F* is the fixed effect of food; and *ε* is the error variance. We estimated broad sense heritability (*H*^2^) by food as *H*^2^ *=* (*σ*^2^*_L_*)/(*σ*^2^*_L_ + σ*^2^*_E_*), where *σ*^2^*_L_* is the variance component among lines, and *σ*^2^*_E_* is the sum of all other sources of variation. In addition, we calculated broadsense heritability across food as *H*^2^ *=* (*σ*^2^*_L_ + σ*^2^*_LF_*)/(*σ*^2^*_L_ + σ*^2^*_LF_ + σ*^2^*_E_*), where *σ*^2^*_LF_* is the line-by-food variance component. We calculated genetic correlations across food *r_NR_ =* (*σ*^2^*_L_)*/(*σ*^2^*_L_ + σ*^2^*_LF_*).

### Genotype-phenotype associations

We performed GWA analyses using line means for all sleep parameters and morphometric measurements using the DGRP pipeline (http://dgrp2.gnets.ncsu.edu/). This pipeline accounts for the effects of *Wolbachia* infection status, major polymorphic inversions and polygenic relatedness ^94^ and implements single-variant tests of association for additive effects of variants with minor allele frequencies ≥ 0.05. We tested the effects of 1,901,174 DNA sequence variants on each trait. We focus our GWA analysis on the associations that represents the interaction of both diets, that is, the difference of phenotypic means between restricted and control food. All annotations that map within or nearby (<± 5,000 bp from gene body) are based on FlyBase release 5.57 (http://www.flybase.org).

### Network analysis

We annotated candidate genes identified by the GWA analyses using FlyBase release 5.57 and mapped gene-gene networks through the genetic interaction database downloaded from FlyBase. We then constructed gene networks using STRING (version 11.0) ^38^ where candidate genes directly interact with each other. We used the following STRING settings: (i) Experiments and Databases as active interaction sources, (ii) high confidence (0.700) as the minimum required interaction score. For network visualization, we used the igraph R package to plot gene networks, where nodes correspond to genes and edges indicate the interaction between genes.

### Gene Ontology analysis

We performed gene ontology (GO) enrichment analysis using PANTHER 17.0 (http://www.pantherdb.org/) ^95^ and STRING 11.0 (https://string-db.org/) ^38^. We used the

DIOPT–*Drosophila* RNAi Screening Center (DRSC) Integrative Ortholog Predictive Tool 9.0, with all available prediction tools and only retrieving the best match when there is more than one match per input gene or protein, to identify human orthologs ^96^.

### Functional analyses

We performed tissue-specific RNAi-mediated knockdown of 17 candidate genes implicated by the GWA analyses using pan-neural *elav-Gal4* and MBs specific *OK107- Gal4* and *201Y-Gal4* drivers (Bloomington, IN, USA) (S17 Table). We crossed males from *UAS*–RNAi lines with virgin females of each of the *Gal4* drivers and reared under standard culture conditions (25 °C, 60–70% humidity,12-hour light:dark cycle) in the two diet conditions. For each cross, we used 35 virgin females and 15 males to obtain comparable levels of offspring density. For the few RNAi lines that yield less offspring in restricted food than in normal food, we used 50 virgin females and 20 males. Adult flies were discarded after three days of egg-laying on NF or RF. After eclosion, adult flies were transferred to vials with NF for 3 days until the sleep behavior or MB morphology was assessed as described above, except that to get more robust results, the sleep data was collected from days 3 to 7 after flies were placed into the DAMs. Sleep measurements were replicated three times for each line. Eight flies of each sex and each diet (NF and RF) were measured in one monitor per replicate. We performed two-way ANOVA followed by Bonferroni’s post hoc test (*p≤ 0.05; **p≤0.01; ***p≤0.001; ****p<0.0001; ns non-significant) to compare the effect of control or candidate gene RNAi knockdown between normal and restricted food. We also performed Welch’s t-test or one-way ANOVA followed by Dunnett’s post hoc test (*p≤ 0.05; **p≤0.01; ***p≤0.001; ****p<0.0001; ns non-significant) to compare the effect of control versus candidate gene RNAi knockdown in each food condition.

## Acknowledgments

We thank Chad A. Highfill and Brandon M. Baker from T.F.C.M. lab (Clemson University) for insightful advice on statistical analyses, Cristian Yáñez from R.A.V. lab for bioinformatic advice, Jimena Sierralta (Universidad de Chile) for reagents and fly stocks, Carlos Oliva (P. Universidad Católica de Chile) for the immunohistochemistry protocol, and Kajan Ratnakumar for critical comments. This work was supported by ANILLO ACT-1401 to P.O. and R.A.V., ICM P09-015F BNI to P.O., UCH-VID INFRA 0440/2018 to R.A.V.

## Author Contributions

G.H.O., P.O. and R.A.V conceived and designed experiments with input from T.F.C.M. F.N.-V. and N.C. maintained fly stocks. G.H.O., F.N.-V., F.V.-M., and N.Z. performed sleep experiments. G.H.O., F.N.-V, F.V.-M., and N.C. performed immunohistochemistry experiments. G.H.O. and F.N.-V. performed morphometric measurements. PO performed adult size experiments. F.N.-V. performed RNAi validation for sleep and morphometric analyses with input from G.H.O. G.H.O. performed statistical analyses with input from R.A.V. G.H.O. and K.O. performed gene networks analyses. G.H.O. and C.M. performed bioinformatic analyses with input from R.A.V. Figures and tables were prepared by G.H.O. The manuscript was written by G.H.O. and P.O. with input from R.A.V. and T.F.C.M.

## Data availability statement

The datasets generated for this study are available on request to the corresponding authors.

## References

1. Bale, T. L., et al. Early life programming and neurodevelopmental disorders. Biol Psychiatry 68, 314–319, doi:10.1016/j.biopsych.2010.05.028 (2010).

2. Li, C., et al. Effect of prenatal and postnatal malnutrition on intellectual functioning in early school-aged children in rural western China. Medicine (Baltimore) 95, e4161, doi:10.1097/MD.0000000000004161 (2016).

3. Keunen, K., van Elburg, R. M., van Bel, F. & Benders, M. J. Impact of nutrition on brain development and its neuroprotective implications following preterm birth. Pediatr Res 77, 148–155, doi:10.1038/pr.2014.171 (2015).

4. Crossland, R. F., et al. Chronic Maternal Low-Protein Diet in Mice Affects Anxiety, Night-Time Energy Expenditure and Sleep Patterns, but Not Circadian Rhythm in Male Offspring. PLoS One 12, e0170127, doi:10.1371/journal.pone.0170127 (2017).

5. Akitake, Y., et al. Moderate maternal food restriction in mice impairs physical growth, behavior, and neurodevelopment of offspring. Nutr Res 35, 76–87, doi:10.1016/j.nutres.2014.10.014 (2015).

6. Kuo, A. H., et al. Maternal nutrient restriction during pregnancy and lactation leads to impaired right ventricular function in young adult baboons. J Physiol 595, 4245–4260, doi:10.1113/JP273928 (2017).

7. Díaz-Cintra, S., Cintra, L., Ortega, A., Kemper, T. & Morgane, P. J. Effects of protein deprivation on pyramidal cells of the visual cortex in rats of three age groups. J Comp Neurol 292, 117–126, doi:10.1002/cne.902920108 (1990).

8. Brown, A. S., van Os, J., Driessens, C., Hoek, H. W. & Susser, E. S. Further evidence of relation between prenatal famine and major affective disorder. Am J Psychiatry 157, 190–195, doi:10.1176/appi.ajp.157.2.190 (2000).

9. Susser, E., et al. Schizophrenia after prenatal famine. Further evidence. Arch Gen Psychiatry 53, 25–31, doi:10.1001/archpsyc.1996.01830010027005 (1996).

10. Hulshoff Pol, H. E., et al. Prenatal exposure to famine and brain morphology in schizophrenia. Am J Psychiatry 157, 1170–1172, doi:10.1176/appi.ajp.157.7.1170 (2000).

11. Falconer, D. S. & Mackay, T. F. C. Introduction to quantitative genetics. 4th edition edn, 464 (Noida : Pearson, 2009).

12. Mackay, T. F. C. & Huang, W. Charting the genotype-phenotype map: lessons from the Drosophila melanogaster Genetic Reference Panel. Wiley Interdiscip Rev Dev Biol 7, doi:10.1002/wdev.289 (2018).

13. Mackay, T. F., et al. The Drosophila melanogaster Genetic Reference Panel. Nature 482, 173–178, doi:10.1038/nature10811 (2012).

14. Unckless, R. L., Rottschaefer, S. M. & Lazzaro, B. P. A genome-wide association study for nutritional indices in Drosophila. G3 (Bethesda) 5, 417–425, doi:10.1534/g3.114.016477 (2015).

15. Jehrke, L., Stewart, F. A., Droste, A. & Beller, M. The impact of genome variation and diet on the metabolic phenotype and microbiome composition of Drosophila melanogaster. Sci Rep 8, 6215, doi:10.1038/s41598-018-24542-5 (2018).

16. Sambandan, D., Carbone, M. A., Anholt, R. R. & Mackay, T. F. Phenotypic plasticity and genotype by environment interaction for olfactory behavior in Drosophila melanogaster. Genetics 179, 1079–1088, doi:10.1534/genetics.108.086769 (2008).

17. Zhou, S., Campbell, T. G., Stone, E. A., Mackay, T. F. & Anholt, R. R. Phenotypic plasticity of the Drosophila transcriptome. PLoS Genet 8, e1002593, doi:10.1371/journal.pgen.1002593 (2012).

18. Reed, L. K., et al. Genotype-by-diet interactions drive metabolic phenotype variation in Drosophila melanogaster. Genetics 185, 1009–1019, doi:10.1534/genetics.109.113571 (2010).

19. Chan, M. S., Chung, K. F., Yung, K. P. & Yeung, W. F. Sleep in schizophrenia: A systematic review and meta-analysis of polysomnographic findings in case-control studies. Sleep Med Rev 32, 69–84, doi:10.1016/j.smrv.2016.03.001 (2017).

20. Wulff, K., Gatti, S., Wettstein, J. G. & Foster, R. G. Sleep and circadian rhythm disruption in psychiatric and neurodegenerative disease. Nat Rev Neurosci 11, 589–599, doi:10.1038/nrn2868 (2010).

21. Cintra, L., Durán, P., Guevara, M. A., Aguilar, A. & Castañón-Cervantes, O. Pre- and post-natal protein malnutrition alters the effect of rapid eye movements sleep-deprivation by the platform-technique upon the electrocorticogram of the circadian sleep-wake cycle and its frequency bands in the rat. Nutr Neurosci 5, 91–101, doi:10.1080/10284150290018964 (2002).

22. Durán, P., Galler, J. R., Cintra, L. & Tonkiss, J. Prenatal malnutrition and sleep states in adult rats: effects of restraint stress. Physiol Behav 89, 156–163, doi:10.1016/j.physbeh.2006.05.045 (2006).

23. Harbison, S. T., McCoy, L. J. & Mackay, T. F. Genome-wide association study of sleep in Drosophila melanogaster. BMC Genomics 14, 281, doi:10.1186/1471-2164-14-281 (2013).

24. Harbison, S. T., et al. Co-regulated transcriptional networks contribute to natural genetic variation in Drosophila sleep. Nat Genet 41, 371–375, doi:10.1038/ng.330 (2009).

25. Pitman, J. L., McGill, J. J., Keegan, K. P. & Allada, R. A dynamic role for the mushroom bodies in promoting sleep in Drosophila. Nature 441, 753–756, doi:10.1038/nature04739 (2006).

26. Joiner, W. J., Crocker, A., White, B. H. & Sehgal, A. Sleep in Drosophila is regulated by adult mushroom bodies. Nature 441, 757–760, doi:10.1038/nature04811 (2006).

27. Sitaraman, D., et al. Propagation of Homeostatic Sleep Signals by Segregated Synaptic Microcircuits of the Drosophila Mushroom Body. Curr Biol 25, 2915–2927, doi:10.1016/j.cub.2015.09.017 (2015).

28. Aso, Y., et al. The neuronal architecture of the mushroom body provides a logic for associative learning. Elife 3, e04577, doi:10.7554/eLife.04577 (2014).

29. Ito, K. & Hotta, Y. Proliferation pattern of postembryonic neuroblasts in the brain of Drosophila melanogaster. Dev Biol 149, 134–148, doi:10.1016/0012-1606(92)90270-q (1992).

30. Lee, T., Lee, A. & Luo, L. Development of the Drosophila mushroom bodies: sequential generation of three distinct types of neurons from a neuroblast. Development 126, 4065–4076 (1999).

31. Lanet, E. & Maurange, C. Building a brain under nutritional restriction: insights on sparing and plasticity from Drosophila studies. Front Physiol 5, 117, doi:10.3389/fphys.2014.00117 (2014).

32. Zwarts, L., et al. The genetic basis of natural variation in mushroom body size in Drosophila melanogaster. Nat Commun 6, 10115, doi:10.1038/ncomms10115 (2015).

33. Vonesch, S. C., Lamparter, D., Mackay, T. F., Bergmann, S. & Hafen, E. Genome-Wide Analysis Reveals Novel Regulators of Growth in Drosophila melanogaster. PLoS Genet 12, e1005616, doi:10.1371/journal.pgen.1005616 (2016).

34. Davie, K., et al. A Single-Cell Transcriptome Atlas of the Aging Drosophila Brain. Cell 174, 982–998.e920, doi:10.1016/j.cell.2018.05.057 (2018).

35. Jenett, A., et al. A GAL4-driver line resource for Drosophila neurobiology. Cell Rep 2, 991–1001, doi:10.1016/j.celrep.2012.09.011 (2012).

36. Li, H. H., et al. A GAL4 driver resource for developmental and behavioral studies on the larval CNS of Drosophila. Cell Rep 8, 897–908, doi:10.1016/j.celrep.2014.06.065 (2014).

37. Shih, M. M., Davis, F. P., Henry, G. L. & Dubnau, J. Nuclear Transcriptomes of the Seven Neuronal Cell Types That Constitute the. G3 (Bethesda) 9, 81–94, doi:10.1534/g3.118.200726 (2019).

38. Szklarczyk, D., et al. STRING v11: protein-protein association networks with increased coverage, supporting functional discovery in genome-wide experimental datasets. Nucleic Acids Res 47, D607–D613, doi:10.1093/nar/gky1131 (2019).

39. Aso, Y., et al. The mushroom body of adult Drosophila characterized by GAL4 drivers. J Neurogenet 23, 156–172, doi:10.1080/01677060802471718 (2009).

40. Luo, L., Liao, Y. J., Jan, L. Y. & Jan, Y. N. Distinct morphogenetic functions of similar small GTPases: Drosophila Drac1 is involved in axonal outgrowth and myoblast fusion. Genes Dev 8, 1787–1802, doi:10.1101/gad.8.15.1787 (1994).

41. DePew, A. T., Aimino, M. A. & Mosca, T. J. The Tenets of Teneurin: Conserved Mechanisms Regulate Diverse Developmental Processes in the. Front Neurosci 13, 27, doi:10.3389/fnins.2019.00027 (2019).

42. Yuan, L. L. & Ganetzky, B. A glial-neuronal signaling pathway revealed by mutations in a neurexin-related protein. Science 283, 1343–1345, doi:10.1126/science.283.5406.1343 (1999).

43. Mathew, D., et al. Recruitment of scribble to the synaptic scaffolding complex requires GUK-holder, a novel DLG binding protein. Curr Biol 12, 531–539, doi:10.1016/s0960-9822(02)00758-3 (2002).

44. Meyer, F. & Moussian, B. Drosophila multiplexin (Dmp) modulates motor axon pathfinding accuracy. Dev Growth Differ 51, 483–498, doi:10.1111/j.1440-169X.2009.01111.x (2009).

45. Wu, C. L., et al. Dock/Nck facilitates PTP61F/PTP1B regulation of insulin signalling. Biochem J 439, 151–159, doi:10.1042/BJ20110799 (2011).

46. Zhang, N., Zhang, J., Cheng, Y. & Howard, K. Identification and genetic analysis of wunen, a gene guiding Drosophila melanogaster germ cell migration. Genetics 143, 1231–1241, doi:10.1093/genetics/143.3.1231 (1996).

47. Shukla, J. P., Deshpande, G. & Shashidhara, L. S. Ataxin 2-binding protein 1 is a context-specific positive regulator of Notch signaling during neurogenesis in. Development 144, 905–915, doi:10.1242/dev.140657 (2017).

48. Bazzi, W., et al. Embryonic hematopoiesis modulates the inflammatory response and larval hematopoiesis in. Elife 7, doi:10.7554/eLife.34890 (2018).

49. Hong, W., Mosca, T. J. & Luo, L. Teneurins instruct synaptic partner matching in an olfactory map. Nature 484, 201–207, doi:10.1038/nature10926 (2012).

50. Morgane, P. J., et al. Prenatal malnutrition and development of the brain. Neurosci Biobehav Rev 17, 91–128, doi:10.1016/s0149-7634(05)80234-9 (1993).

51. Li, Y., Zhao, L., Yu, D. & Ding, G. Exposure to the Chinese famine in early life and depression in adulthood. Psychol Health Med 23, 952–957, doi:10.1080/13548506.2018.1434314 (2018).

52. He, P., et al. Prenatal malnutrition and adult cognitive impairment: a natural experiment from the 1959-1961 Chinese famine. Br J Nutr 120, 198–203, doi:10.1017/S0007114518000958 (2018).

53. Thurmond, J., et al. FlyBase 2.0: the next generation. Nucleic Acids Res 47, D759–D765, doi:10.1093/nar/gky1003 (2019).

54. King, E. G. & Long, A. D. The Beavis Effect in Next-Generation Mapping Panels in. G3 (Bethesda) 7, 1643–1652, doi:10.1534/g3.117.041426 (2017).

55. Mottus, R. C., et al. Unique gene organization: alternative splicing in Drosophila produces two structurally unrelated proteins. Gene 198, 229–236, doi:10.1016/s0378-1119(97)00319-3 (1997).

56. Cabrera y Poch, H. L., Arribas, C. & Izquierdo, M. Sequence of a Drosophila cDNA encoding a ubiquitin gene fusion to a 52-aa ribosomal protein tail. Nucleic Acids Res 18, 3994, doi:10.1093/nar/18.13.3994 (1990).

57. Lambertsson, A. The minute genes in Drosophila and their molecular functions. Adv Genet 38, 69–134, doi:10.1016/s0065-2660(08)60142-x (1998).

58. Marygold, S. J., et al. The ribosomal protein genes and Minute loci of Drosophila melanogaster. Genome Biol 8, R216, doi:10.1186/gb-2007-8-10-r216 (2007).

59. Hasygar, K. & Hietakangas, V. p53- and ERK7-dependent ribosome surveillance response regulates Drosophila insulin-like peptide secretion. PLoS Genet 10, e1004764, doi:10.1371/journal.pgen.1004764 (2014).

60. Fisher, C. E. & Howie, S. E. The role of megalin (LRP-2/Gp330) during development. Dev Biol 296, 279–297, doi:10.1016/j.ydbio.2006.06.007 (2006).

61. Huang, R., et al. High-fat diet enhances starvation-induced hyperactivity via sensitizing hunger-sensing neurons in. Elife 9, doi:10.7554/eLife.53103 (2020).

62. Desai, C. J., Krueger, N. X., Saito, H. & Zinn, K. Competition and cooperation among receptor tyrosine phosphatases control motoneuron growth cone guidance in Drosophila. Development 124, 1941–1952 (1997).

63. Goossens, T., et al. The Drosophila L1CAM homolog Neuroglian signals through distinct pathways to control different aspects of mushroom body axon development. Development 138, 1595–1605, doi:10.1242/dev.052787 (2011).

64. Lin, D. M., Fetter, R. D., Kopczynski, C., Grenningloh, G. & Goodman, C. S. Genetic analysis of Fasciclin II in Drosophila: defasciculation, refasciculation, and altered fasciculation. Neuron 13, 1055–1069, doi:10.1016/0896-6273(94)90045-0 (1994).

65. Ashley, J., Packard, M., Ataman, B. & Budnik, V. Fasciclin II signals new synapse formation through amyloid precursor protein and the scaffolding protein dX11/Mint. J Neurosci 25, 5943–5955, doi:10.1523/JNEUROSCI.1144-05.2005 (2005).

66. Winkle, C. C. et al. Trim9 Deletion Alters the Morphogenesis of Developing and Adult-Born Hippocampal Neurons and Impairs Spatial Learning and Memory. J Neurosci 36, 4940–4958, doi:10.1523/JNEUROSCI.3876-15.2016 (2016).

67. Dashti, H. S., et al. Genome-wide association study identifies genetic loci for self-reported habitual sleep duration supported by accelerometer-derived estimates. Nat Commun 10, 1100, doi:10.1038/s41467-019-08917-4 (2019).

68. Jansen, P. R., et al. Genome-wide analysis of insomnia in 1,331,010 individuals identifies new risk loci and functional pathways. Nat Genet 51, 394–403, doi:10.1038/s41588-018-0333-3 (2019).

69. Lane, J. M., et al. Biological and clinical insights from genetics of insomnia symptoms. Nat Genet 51, 387–393, doi:10.1038/s41588-019-0361-7 (2019).

70. Jones, S. E., et al. Genetic studies of accelerometer-based sleep measures yield new insights into human sleep behaviour. Nat Commun 10, 1585, doi:10.1038/s41467-019-09576-1 (2019).

71. Denault, J. B., et al. Processing of proendothelin-1 by human furin convertase. FEBS Lett 362, 276–280, doi:10.1016/0014-5793(95)00249-9 (1995).

72. Seidah, N. G., Benjannet, S., Pareek, S., Chrétien, M. & Murphy, R. A. Cellular processing of the neurotrophin precursors of NT3 and BDNF by the mammalian proprotein convertases. FEBS Lett 379, 247–250, doi:10.1016/0014-5793(95)01520-5 (1996).

73. Faraguna, U., Vyazovskiy, V. V., Nelson, A. B., Tononi, G. & Cirelli, C. A causal role for brain-derived neurotrophic factor in the homeostatic regulation of sleep. J Neurosci 28, 4088–4095, doi:10.1523/JNEUROSCI.5510-07.2008 (2008).

74. Huang, E. J. & Reichardt, L. F. Neurotrophins: roles in neuronal development and function. Annu Rev Neurosci 24, 677–736, doi:10.1146/annurev.neuro.24.1.677 (2001).

75. McLaughlin, S. & Dixon, J. E. Alternative splicing gives rise to a nuclear protein tyrosine phosphatase in Drosophila. J Biol Chem 268, 6839–6842 (1993).

76. Nakahata, S. & Kawamoto, S. Tissue-dependent isoforms of mammalian Fox-1 homologs are associated with tissue-specific splicing activities. Nucleic Acids Res 33, 2078–2089, doi:10.1093/nar/gki338 (2005).

77. Chell, J. M. & Brand, A. H. Nutrition-responsive glia control exit of neural stem cells from quiescence. Cell 143, 1161–1173, doi:10.1016/j.cell.2010.12.007 (2010).

78. Carreira-Rosario, A., et al. Repression of Pumilio Protein Expression by Rbfox1 Promotes Germ Cell Differentiation. Dev Cell 36, 562–571, doi:10.1016/j.devcel.2016.02.010 (2016).

79. Lee, J. A., et al. Cytoplasmic Rbfox1 Regulates the Expression of Synaptic and Autism-Related Genes. Neuron 89, 113–128, doi:10.1016/j.neuron.2015.11.025 (2016).

80. Weyn-Vanhentenryck, S. M., et al. HITS-CLIP and integrative modeling define the Rbfox splicing-regulatory network linked to brain development and autism. Cell Rep 6, 1139–1152, doi:10.1016/j.celrep.2014.02.005 (2014).

81. Potter, C. J., Huang, H. & Xu, T. Drosophila Tsc1 functions with Tsc2 to antagonize insulin signaling in regulating cell growth, cell proliferation, and organ size. Cell 105, 357–368, doi:10.1016/s0092-8674(01)00333-6 (2001).

82. Gu, T., Zhao, T. & Hewes, R. S. Insulin signaling regulates neurite growth during metamorphic neuronal remodeling. Biol Open 3, 81–93, doi:10.1242/bio.20136437 (2014).

83. Gehman, L. T., et al. The splicing regulator Rbfox1 (A2BP1) controls neuronal excitation in the mammalian brain. Nat Genet 43, 706–711, doi:10.1038/ng.841 (2011).

84. Gehman, L. T., et al. The splicing regulator Rbfox2 is required for both cerebellar development and mature motor function. Genes Dev 26, 445–460, doi:10.1101/gad.182477.111 (2012).

85. Reed, L. K., et al. Systems genomics of metabolic phenotypes in wild-type Drosophila melanogaster. Genetics 197, 781–793, doi:10.1534/genetics.114.163857 (2014).

86. Mohr, S. E., et al. Methods and tools for spatial mapping of single-cell RNAseq clusters in Drosophila. Genetics 217, doi:10.1093/genetics/iyab019 (2021).

87. Bass, T. M., et al. Optimization of dietary restriction protocols in Drosophila. J Gerontol A Biol Sci Med Sci 62, 1071–1081, doi:10.1093/gerona/62.10.1071 (2007).

88. Medina-Yáñez, I., Olivares, G. H., Vega-Macaya, F., Mlodzik, M. & Olguín, P. Phosphatidic acid increases Notch signalling by affecting Sanpodo trafficking during Drosophila sensory organ development. Sci Rep 10, 21731, doi:10.1038/s41598-020-78831-z (2020).

89. Shaw, P. J., Cirelli, C., Greenspan, R. J. & Tononi, G. Correlates of sleep and waking in Drosophila melanogaster. Science 287, 1834–1837, doi:10.1126/science.287.5459.1834 (2000).

90. Hendricks, J. C., et al. Rest in Drosophila is a sleep-like state. Neuron 25, 129–138, doi:10.1016/s0896-6273(00)80877-6 (2000).

91. Gilestro, G. F. & Cirelli, C. pySolo: a complete suite for sleep analysis in Drosophila. Bioinformatics 25, 1466–1467, doi:10.1093/bioinformatics/btp237 (2009).

92. Rollmann, S. M., et al. Pleiotropic effects of Drosophila neuralized on complex behaviors and brain structure. Genetics 179, 1327–1336, doi:10.1534/genetics.108.088435 (2008).

93. Schindelin, J., et al. Fiji: an open-source platform for biological-image analysis. Nat Methods 9, 676–682, doi:10.1038/nmeth.2019 (2012).

94. Huang, W., et al. Natural variation in genome architecture among 205 Drosophila melanogaster Genetic Reference Panel lines. Genome Res 24, 1193–1208, doi:10.1101/gr.171546.113 (2014).

95. Mi, H., Muruganujan, A., Ebert, D., Huang, X. & Thomas, P. D. PANTHER version 14: more genomes, a new PANTHER GO-slim and improvements in enrichment analysis tools. Nucleic Acids Res 47, D419–D426, doi:10.1093/nar/gky1038 (2019).

96. Rotival, M., et al. Integrating genome-wide genetic variations and monocyte expression data reveals trans-regulated gene modules in humans. PLoS Genet 7, e1002367, doi:10.1371/journal.pgen.1002367 (2011).

