## Supplementary Information for "Early-life nutrition interacts with developmental genes to shape the brain and sleep behavior in *Drosophila melanogaster*"

### Supporting Information

**S1 Fig. Body size response to early-life nutrition.**

(A) Scheme showing body size measurements. Box plots of Oregon-R body size measurements of (B, E) interocular distance (IOD), (C, F) notum length (NL) and (D, G) IOD/NL ratio from females (B-D) or males (E-G) reared under Normal (blue) or Restricted (red) food. We performed a two-way non-parametric t-Test followed by Mann-Whitney's post hoc test (**p*≤ 0.05; ***p*≤0.01; ****p*≤0.001; *****p*<0.0001; ns non-significant) to compare the body size traits between normal and restricted food. (B-D) females n = 50 for N and n = 40 for R; (E-G) males n = 54 for N and n = 39 for R. a.u.: arbitrary units; N: Normal food; R: Restricted food.

**S2 Fig. Sleep trait response to early-life nutrition in females.**

Histograms of female sleep trait mean + SEM for (A) Day sleep, (B) Day average bout length, (C) Day bout number, (D) Night average bout length, (E) Night bout number, (F) Total sleep, (G) Latency, and (H) Waking activity. Reaction norms for sleep traits. (A') Day sleep, (B') Day average bout length, (C') Day bout number, (D') Night average bout length, (E') Night bout number, (F') Total sleep, (G') Latency, and (H') Waking activity. Each DGRP line is represented by a different color. N: Normal food; R: Restricted food.

**S3 Fig. Sleep trait response to early-life nutrition in males.**

Histograms of male sleep trait mean + SEM for (A) Day sleep, (B) Day average bout length, (C) Day bout number, (D) Night average bout length, (E) Night bout number, (F) Total sleep, (G) Latency, and (H) Waking activity. Reaction norms for sleep traits. (A')

Day sleep, (B') Day average bout length, (C') Day bout number, (D') Night average bout length, (E') Night bout number, (F') Total sleep, (G') Latency, and (H') Waking activity. Each DGRP line is represented by a different color. N: Normal food; R: Restricted food.

**S4 Fig. Q-Q plot of *P*-values from the DGRP single variant GWA analysis of sleep traits in response to early-life nutrition in females.** (A) Day sleep, (B) Day bout number, (C) Night sleep, (D) Night average bout length, (E) Night bout number, (F) Total sleep, (G) Latency, and (H) Waking activity.

**S5 Fig. Q-Q plot of *P*-values from the DGRP single variant GWA analysis of sleep traits in response to early-life nutrition in males.** (A) Day sleep, (B) Day bout number, (C) Night sleep, (D) Night average bout length, (E) Night bout number, (F) Total sleep, (G) Latency, and (H) Waking activity.

**S6 Fig. Q-Q plot of *P*-values from the DGRP single variant GWA analysis of MBs morphology traits in response to early-life nutrition in females.** (A) $\alpha$-lobe length, (B) $\alpha$-lobe width, (C) $\beta$-lobe length, and (D) $\beta$-lobe width.

**S7 Fig. Genome-wide association analyses for sleep traits in response to early-life nutrition in females.**

Manhattan plots of all SNPs associated with female sleep traits difference between diets. (A) Day sleep, (B) Night sleep, (C) Night average bout length, and (D) Latency.

The green line indicates the nominal *P*-value ≤ $10^{-5}$ reporting threshold. *P*-values are plotted as $-\log_{10}$(*P*-value).

**S8 Fig. Genome-wide association analyses for sleep traits in response to early-life nutrition in males.**

Manhattan plots of all SNPs associated with male sleep traits difference between diets. (A) Day bout number, (B) Night sleep, (C) Night average bout length, (D) Night bout number, and (E) Latency. The green line indicates the nominal *P*-value ≤ $10^{-5}$ reporting threshold. *P*-values are plotted as $-\log_{10}$(*P*-value).

**S9 Fig. Genomic locations of variants associated with sleep traits.**

Variants were annotated as intergenic (outside the transcribed region of an annotated gene), UTR (5' and 3'), coding or intron according to the site class. (A) Genomic localization of variants as percentage (%) of sleep associated variants in both sexes for all sleep traits. (B) Genomic localization of variants as percentage (%) of sleep associated variants in females for each trait. (C) Genomic localization of variants as percentage (%) of sleep associated variants in males for each trait.

**S10 Fig. Gene Ontology enrichment analysis of candidate genes associated with sleep traits.**

Biological process enrichment analysis of candidate sleep genes. Fisher's exact test, Bonferroni's correction for multiple testing *P*-value. *P*-values are $-\log_{10}$ transformed.

**S11 Fig. Human orthologs of sleep associated genes and their role in complex traits and disease.**

(A) Human orthologs associated with sleep phenotypes. (B) Human orthologs associated with brain structure phenotypes.

**Fig 12. RNAi-mediated knockdown of candidate genes associated with GENI in sleep affects sleep response to early-life nutrition.**

(A-D) Sleep analyses of flies expressing candidate genes RNAis in the MBs or in all neurons in NF and RF. Reaction norms depict sleep trait mean in the two experimental conditions: RNAi control (dashed lines) and RNAi candidate gene (solid lines) using a pan-neural driver (*elav-Gal4*, orange lines) or MBs driver (*OK107-Gal4*, blue lines) from flies reared under Normal or Restricted food. We performed two-way ANOVA followed by Bonferroni's post hoc test (*p≤ 0.05; **p≤0.01; ***p≤0.001; ****p<0.0001; ns non-significant) to compare the effect of control or candidate gene RNAi knockdown between normal and restricted food (see color coded asterisks and ns). We also performed one-way ANOVA followed by Dunnett's post hoc test (*p≤ 0.05; **p≤0.01; ***p≤0.001; ****p<0.0001; ns non-significant) to compare the effect of control versus candidate gene RNAi knockdown in each food condition (see gray coded asterisks and ns). The candidate's gene name and sex of adult flies are indicated at the top of each plot. N: Normal food; R: Restricted food.

**S13 Fig. Summary table of the RNAi-mediated knockdown of the sleep associated genes in response to early-life nutrition.**

Sleep analyses in response to early-life nutrition when specific genes are knockdown at all neurons with *elav-Gal4* driver (orange-colored columns) or at MBs with *OK107-Gal4* driver (light-blue-colored columns) in females (A) and males (B). Results were summarized according to their comparison with their respective genetic background control RNAi response to normal vs. restricted diet. (-): no change in the sleep trait response to nutritional restriction compared with the response of control RNAi; (↑) or (↓): increased or decreased, respectively, sleep trait response to nutritional restriction compared with the response of control RNAi. Source traits from which the SNPs were significatively associated in GWAS are indicated for each gene.

**S14 Fig. RNAi-mediated knockdown of candidate genes associated with GENI in sleep affects sleep response to early-life nutrition.**

(A-D) Sleep analyses in response to early-life nutrition when specific candidate genes are knockdown with a different RNAi (denoted as "(2)") at MBs with *OK107-Gal4* or *201Y-Gal4* drivers. Reaction norms depict sleep trait mean in the two experimental conditions: RNAi control (dashed lines) and RNAi candidate gene (solid lines) using MBs drivers (*OK107-Gal4*, blue lines; *201Y-Gal4*, green lines) from flies reared under Normal or Restricted food. We performed two-way ANOVA followed by Bonferroni's post hoc test (*p≤ 0.05; **p≤0.01; ***p≤0.001; ****p<0.0001; ns non-significant) to compare the effect of control or candidate gene RNAi knockdown between normal and restricted food (see color coded asterisks and ns). We also performed one-way ANOVA followed by Dunnett's post hoc test (*p≤ 0.05; **p≤0.01; ***p≤0.001; ****p<0.0001; ns non-significant) to compare the effect of control versus candidate gene RNAi knockdown in each food condition (see gray coded asterisks and ns). The candidate's gene name and sex of adult flies are indicated at the top of each plot. N: Normal food; R: Restricted food.

**S15 Fig. Summary table of the RNAi-mediated knockdown of the MBs morphology associated genes in response to early-life nutrition.**

Sleep analyses in response to early-life nutrition when specific genes are knockdown at all neurons with *elav-Gal4* driver (orange-colored columns) or at MBs with *OK107-Gal4* driver (blue-colored columns) or *201Y-Gal4* driver (green-colored columns) in females (A) and males (B). Results were summarized according to their comparison with their respective genetic background control RNAi response to normal vs. restricted diet. (-): no change in the sleep trait response to nutritional restriction compared with the response of control RNAi; (↑) or (↓): increased or decreased, respectively, sleep trait response to nutritional restriction compared with the response of control RNAi. ND: not determined. Note that a different RNAi (denoted as "(2)") was also evaluated. Source traits from which the SNPs were significatively associated in GWAS are indicated for each gene.

**S16 Fig. RNAi-mediated knockdown of candidate genes associated with GENI in sleep and morphology affects MBs morphology in response to early-life nutrition.**

MBs morphology analyses in response to early-life nutrition after knockdown of candidate genes in the MBs. (A-L) Reaction norms depict female MBs morphology traits comparing two experimental conditions: RNAi control (dashed lines) and RNAi candidate gene (solid lines) using a MBs driver *OK107-Gal4* from flies reared under Normal or Restricted food. We performed (A-D) Welch's t-test or (E-L) two-way ANOVA followed by Bonferroni's post hoc test (*p≤ 0.05; **p≤0.01; ***p≤0.001; ****p<0.0001; ns non-significant) to compare the effect of control or candidate gene RNAi knockdown between normal and restricted food (see color coded asterisks and ns). We also performed one-way ANOVA followed by Dunnett's post hoc test (*p≤ 0.05; **p≤0.01; ***p≤0.001; ****p<0.0001; ns non-significant) to compare the effect of control versus candidate gene RNAi knockdown in each food condition (see gray coded asterisks and ns). The candidate's gene name and sex of adult flies are indicated at the top of each plot. N: Normal food; R: Restricted food.

**S17 Fig. Summary table of the RNAi-mediated knockdown of sleep associated genes results in altered MBs morphometric measurements in response to early-life nutrition.**

MBs morphology traits analyses in response to early-life nutrition when specific genes are knockdown at MBs with *OK107-Gal4* driver in females. Results were summarized according to their comparison with their respective genetic background control RNAi response to normal vs. restricted diet. (-): no change in the MB morphology trait response to nutritional restriction compared with the response of control RNAi; (↑) or (↓): increased or decreased, respectively, MB morphology trait response to nutritional restriction compared with the response of control RNAi.

**S1 Table. DGRP raw sleep data.**

F: female; M: male; N: normal; R: restricted. (XLSX).

**S2 Table. DGRP line means for all sleep traits.**

(A) Sleep traits from females reared on Normal food. (B) Sleep traits from females reared on Restricted food. (C) Sleep traits from males reared on Normal food. (D) Sleep traits from males reared on Restricted food. F: female; M: male; N: normal; R: restricted. (XLSX).

**S3 Table. Analyses of variance of sleep traits.**

Food, Sex, and their interaction are fixed effects, the rest are random. Mixed model, two-way factorial ANOVAs are given for males and females as well as reduced models by Sex and Food. L: DGRP Line; R: Replicate; S: Sex; F: Food; df: degrees of freedom; MS: Type III mean squares; F: F-ratio test; *P*: *P*-value; $\sigma^2$: variance component estimate; *SE*: standard error; $H^2$: Broad-sense heritability; *r*: cross-food, cross-sex genetic correlations; $i^2$: interaction coefficient across food. (A) Day sleep, (B) Day average bout length, (C) Day bout number, (D) Night sleep, (E) Night average bout length, (F) Night bout number, (G) Total sleep, (H) Latency, (I) Waking activity. (XLSX).

**S4 Table. Percentage of gross MB phenotypes observed in the 40 DGRP lines.**

(A) Phenotypes in Normal food. (B) Phenotypes in Restricted Food. (XLSX).

**S5 Table. DGRP raw MBs data.**

F: female; N: normal; R: restricted. Normalized means are given in arbitrary units (a.u.). (XLSX).

**S6 Table. DGRP line means for all MBs traits.**

(A) MBs traits from females reared on Normal food. (B) MBs traits from females reared on Restricted food. Normalized means are given in arbitrary units (a.u.). F: female; M: male; N: normal; R: restricted. (XLSX).

**S7 Table. Analyses of variance of MBs traits.**

Food, and its interaction are fixed effects, the rest are random. Mixed model, one-way factorial ANOVAs are given females as well as reduced models by Food. L: DGRP Line; F: Food; df: degrees of freedom; MS: Type III mean squares; F: F-ratio test; *P*: *P*-value; $\sigma^2$: variance component estimate; *SE*: standard error; $H^2$: Broad-sense heritability; *r*: cross-food genetic correlation; $i^2$: interaction coefficient across food. (A) $\alpha$-lobe length, (B) $\alpha$-lobe width, (C) $\beta$-lobe length, (D) $\beta$-lobe width. (XLSX).

**S8 Table. Results of genome wide association (GWA) analyses for sleep behavior.**

(A) Top variants ($P \leq 1 \times 10^{-5}$) and associated genes for each sleep trait in females. (B) Variants and genes for the sleep traits in females. (C) Top variants ($P \leq 1 \times 10^{-5}$) and associated genes for each sleep trait in males. (D) Variants and genes for the sleep traits in males. (E) Genes for the sleep traits in females and males. (F) GWAS summary for sleep behavior. (G) Gene ontology enrichment analysis for the sleep GWA analyses (PANTHER). (H) Genes for the sleep traits associated with Nervous System

Development Gene Ontology (GO:0007399). (I) Human orthologs of sleep associated genes, indicating relevant disease and traits (DIOPT-Disease). (XLSX).

**S9 Table. Comparison of sleep associated genes with genes expressed in MBs from previous reports.**

(A) A single-cell atlas of adult fly brains [90] Table S3 - Marker genes and statistics. (B) Adult brain MB expressed GAL4 lines [35] FlyLight web server. (C) Third instar larva brain MB expressed GAL4 lines [36] FlyLight web server. (D) Genes enriched in individual MB classes [91] Table S2. (XLSX).

**S10 Table. Sleep data for RNAi and control genotypes.**

(A) Raw sleep data of RNAi knockdown. (B) Summary data, mean and SEM for each RNAi in each sleep parameter in females. RNAi lines were driven to all neurons with *elav-Gal4*. (C) Summary data, mean and SEM for each RNAi in each sleep parameter in males. RNAi lines were driven to all neurons with *elav-Gal4*. (D) Summary data, mean and SEM for each RNAi in each sleep parameter in females. RNAi lines were driven to MBs with *OK107-Gal4*. (E) Summary data, mean and SEM for each RNAi in each sleep parameter in males. RNAi lines were driven to MBs with *OK107-Gal4*. (F) Summary data, mean and SEM for each RNAi in each sleep parameter in females. RNAi lines were driven to MBs with *201Y-Gal4*. (G) Summary data, mean and SEM for each RNAi in each sleep parameter in males. RNAi lines were driven to MBs with *201Y-Gal4*. We performed by two-way ANOVA followed by Bonferroni's post hoc test (*$p \leq 0.05$; **$p \leq 0.01$; ***$p \leq 0.001$; ****$p < 0.0001$; ns non-significant) to compare the effect of

RNAi knockdown between normal and restricted food considering their appropriate genetic background control RNAi. F: female; M: male; N: normal; R: restricted; ND: not determined. Note that a different RNAi (denoted as “(2)”) was also evaluated for a subset of genes. (XLSX).

**S11 Table. MB morphometric data for RNAi and control genotypes.**

(A) Raw normalized MB morphometric data of RNAi knockdown. (B) Summary data with mean and SEM for each RNAi in each morphology parameter. RNAi lines were driven to MBs with *OK107-Gal4*. Normalized means are given in arbitrary units (a.u.). Two-way ANOVA followed by Bonferroni’s post hoc test (*$p \leq 0.05$; **$p \leq 0.01$; ***$p \leq 0.001$; ****$p < 0.0001$; ns non-significant) to compare the effect of control or candidate gene RNAi knockdown between normal and restricted food. F: female; M: male; N: normal; R: restricted. (XLSX).

**S12 Table. Comparison of genes associated with sleep behavior from previous reports.**

(A) Genes associated with natural variation of sleep [23] Additional file 9. (B) Genes associated with natural variation of sleep [23] Additional file 10. (C) Genes associated with natural variation of sleep [24] Supplementary Table 2a. (XLSX).

**S13 Table. *Drosophila* lines used in this study.**

(A) DGRP lines for sleep behavior. (B) DGRP lines for MBs morphology. (C) RNAi lines, control genotypes and GAL4 driver line. Note that a different RNAi line for the same gene is designated as "(2)". (XLSX).

**S14 Table. Comparison of genes associated with sleep behavior in humans and human orthologs of *Drosophila* genes associated with sleep response to early-life nutrition.** (A) Genes associated with sleep duration and diseases (Dashti et al., 2019) [70] Supplementary Data 1. Genome-wide association signals ($P < 5\text{x}10^{-8}$) for sleep duration. (B) Genes associated with sleep duration and diseases (Dashti et al., 2019) [70] Supplementary Data 4. Genome-wide association signals ($P < 5\text{x}10^{-8}$) for short sleep. (C) Genes associated with sleep duration and diseases (Dashti et al., 2019) [70] Supplementary Data 7. Additional new loci ($P < 5\text{x}10^{-8}$) unraveled for sleep duration. (D) Genes associated with sleep duration and diseases (Dashti et al., 2019) [70] Supplementary Data 8. Genetic variants for self-reported sleep duration association. (E) Genes associated with sleep duration and diseases (Dashti et al., 2019) [70] Supplementary Data 9. Genetic variants for self-reported short and long sleep associations. (F) Genes associated with sleep duration and diseases (Dashti et al., 2019) [70] Supplementary Data 10. Annotation of genes under association signals for sleep duration. (G) Genes associated with self-reported habitual sleep duration (Jones et al., 2019) [73] Table 3 Summary statistics for 47 genetic associations. (H) Genes associated with self-reported insomnia symptoms (Lane et al., 2019) [72] Supplementary Table 2. Genome-wide significant loci ($P < 5\text{x}10^{-8}$) associated with insomnia symptoms. (I) Genes associated with insomnia (Jansen et al., 2019) [71]

Supplementary Table 5. Overview of gene-mapping of insomnia by four methods for males and females in gender-specific analysis. (J) Summary of all common genes (A to I). (XLSX).
