## Supplementary Figures for "Early-life nutrition interacts with developmental genes to shape the brain and sleep behavior in *Drosophila melanogaster*"

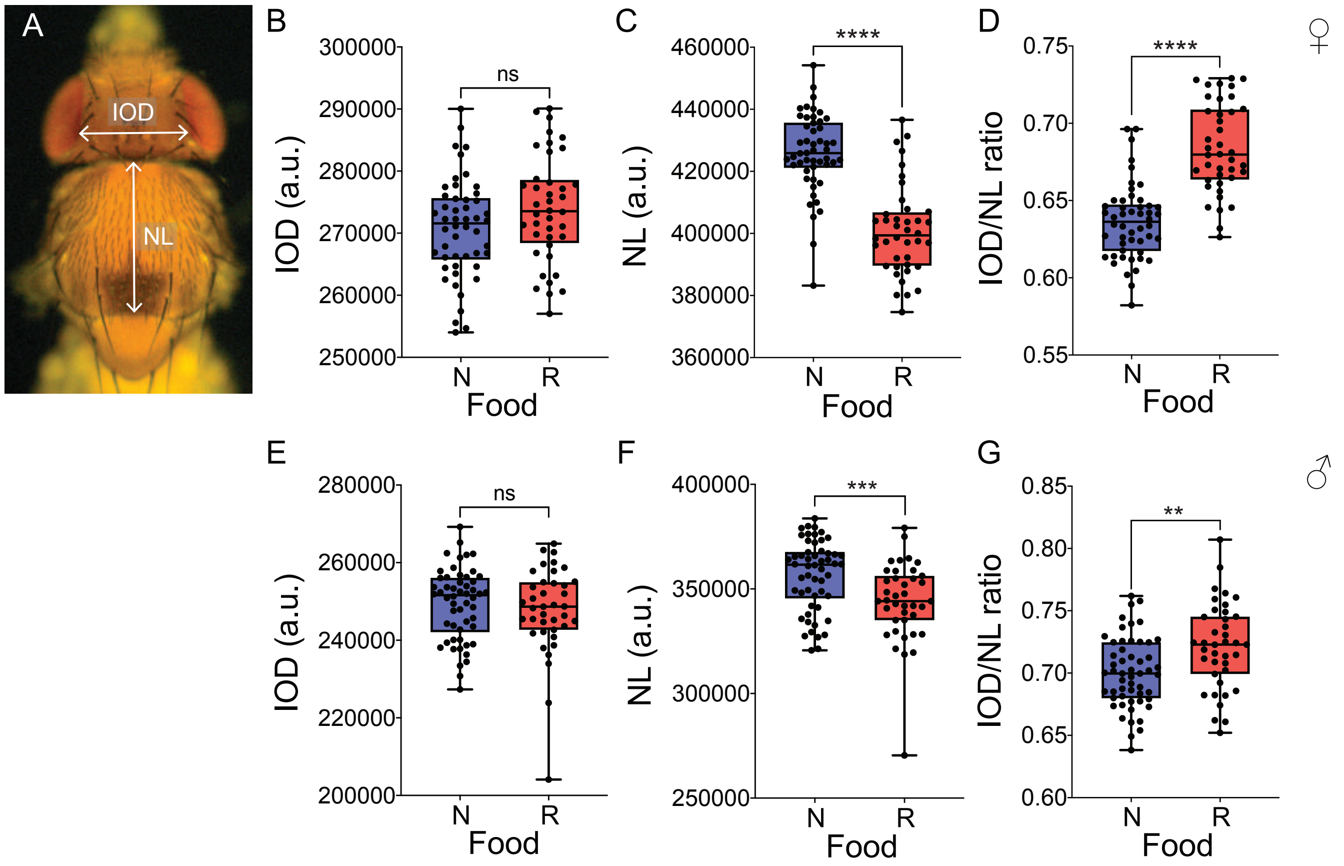


S1 Fig. Olivares et al., 2022.

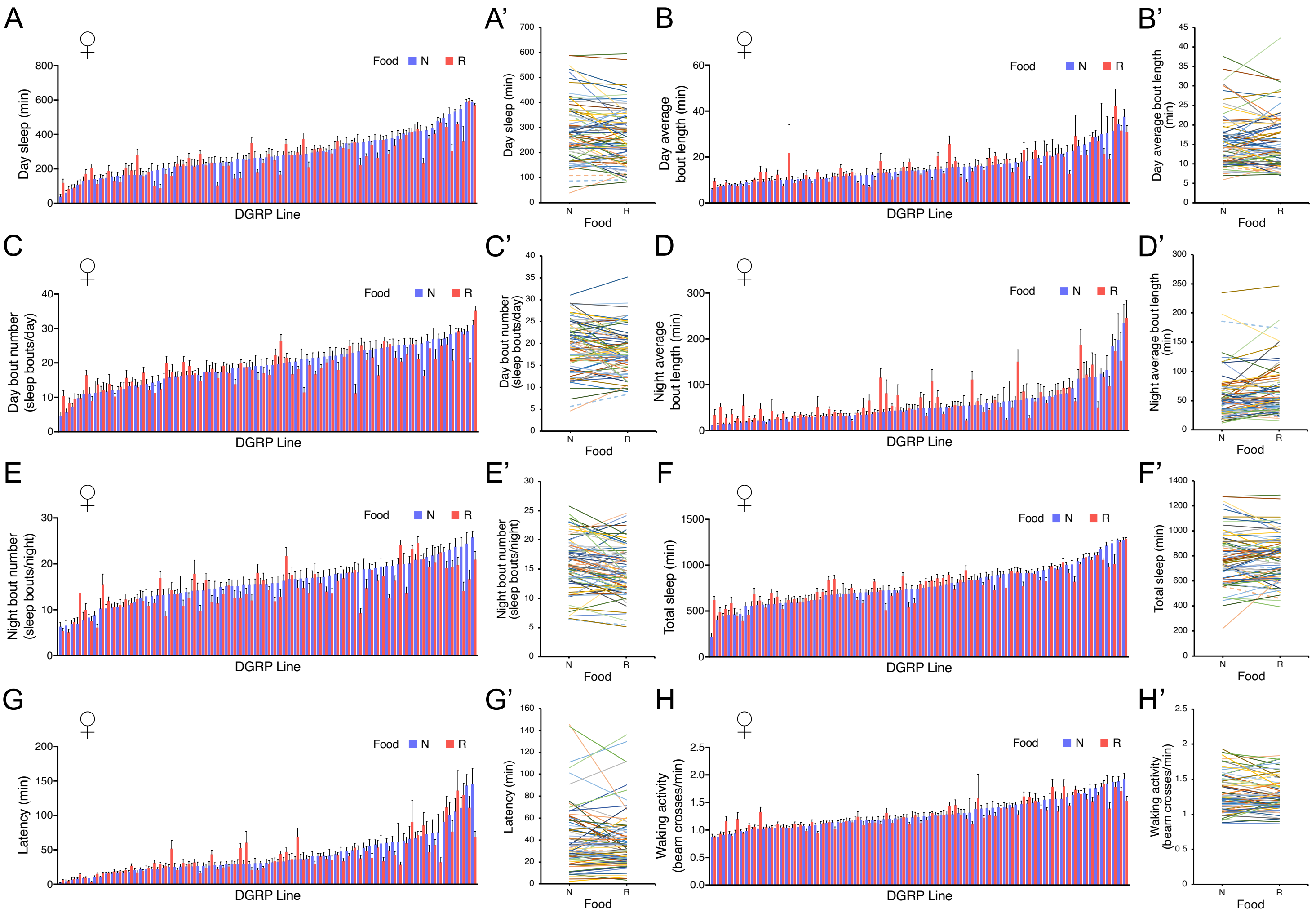


S2 Fig. Olivares et al., 2022.

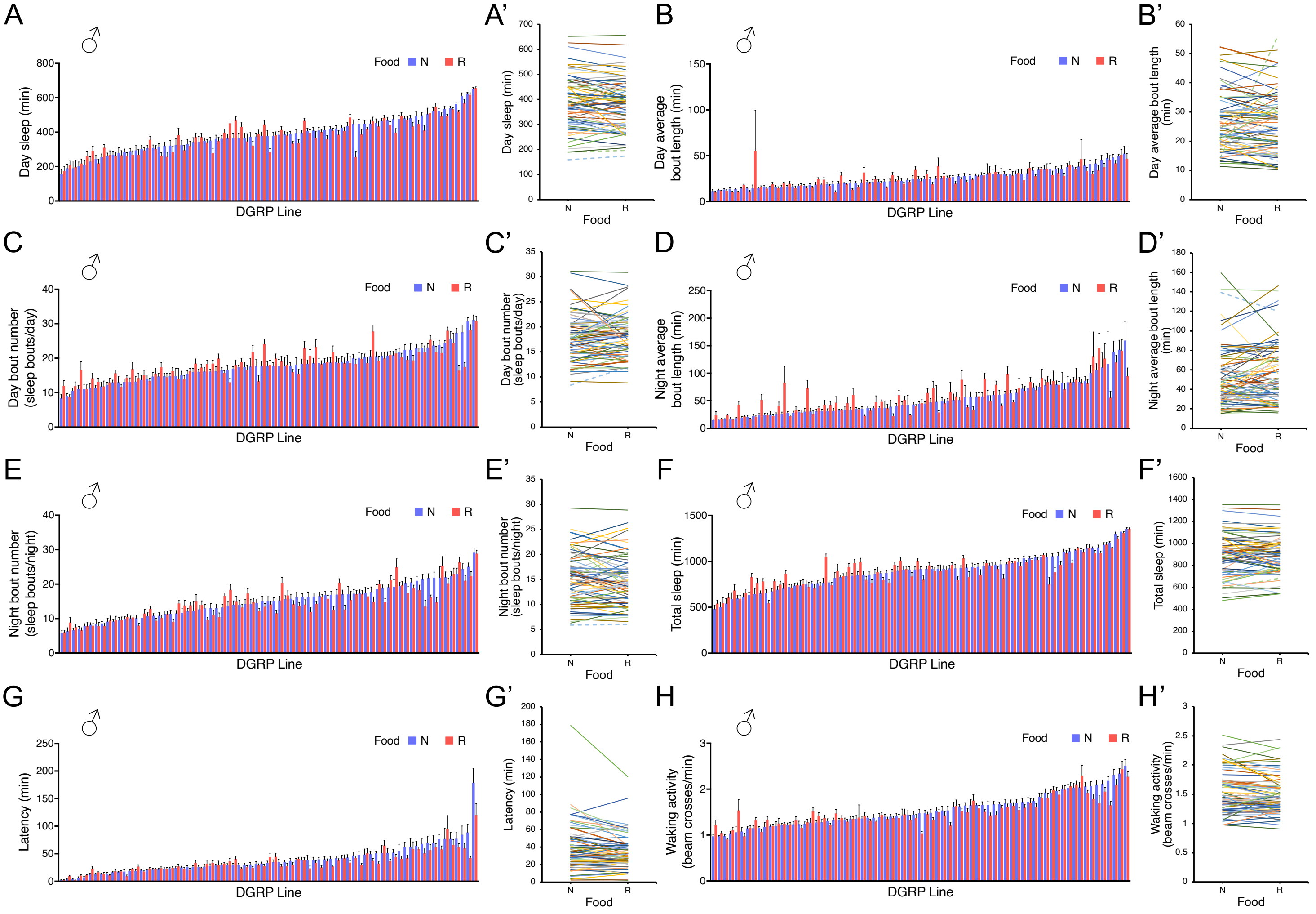


S3 Fig. Olivares et al., 2022.

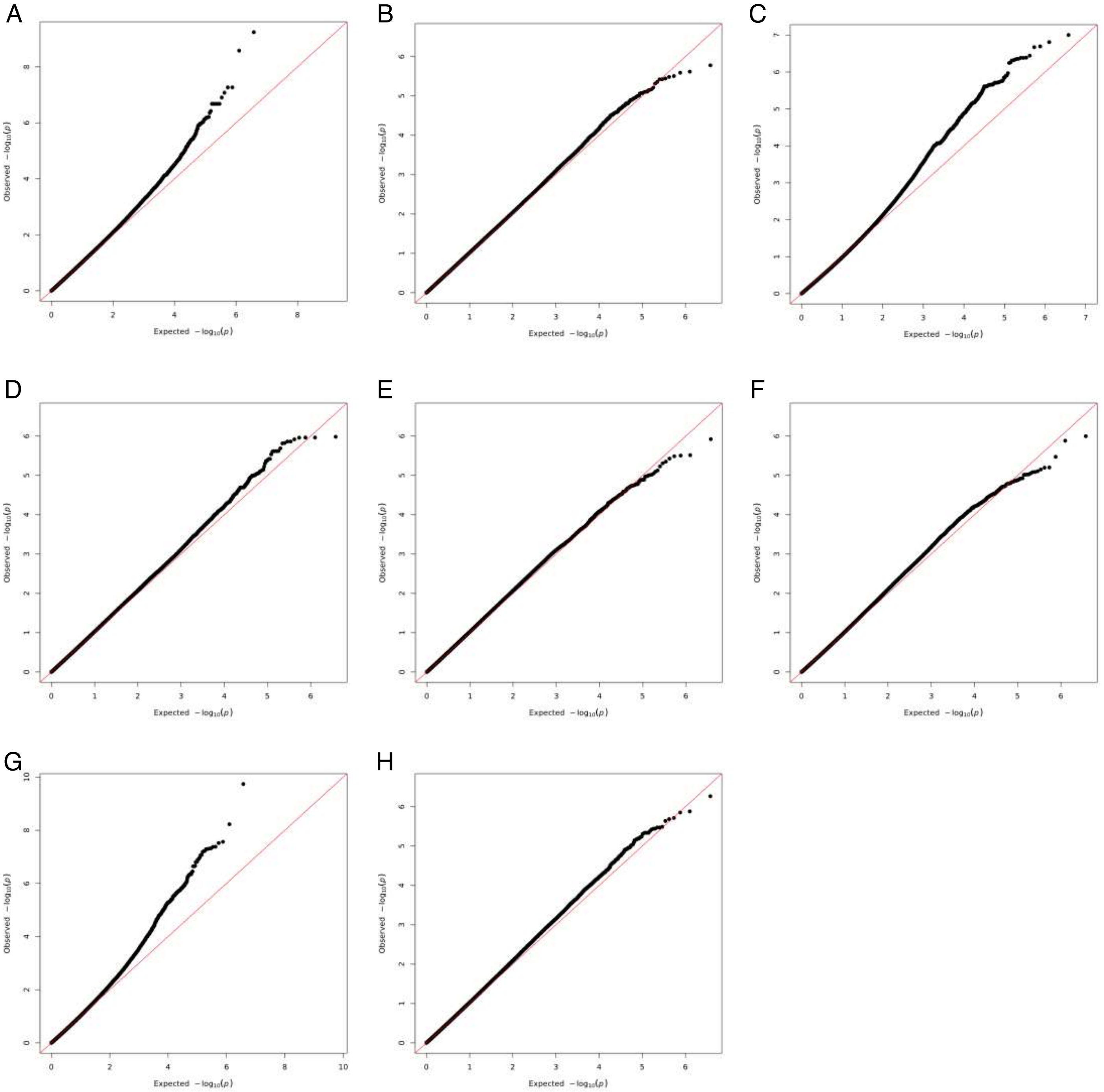


S4 Fig. Olivares et al., 2022.

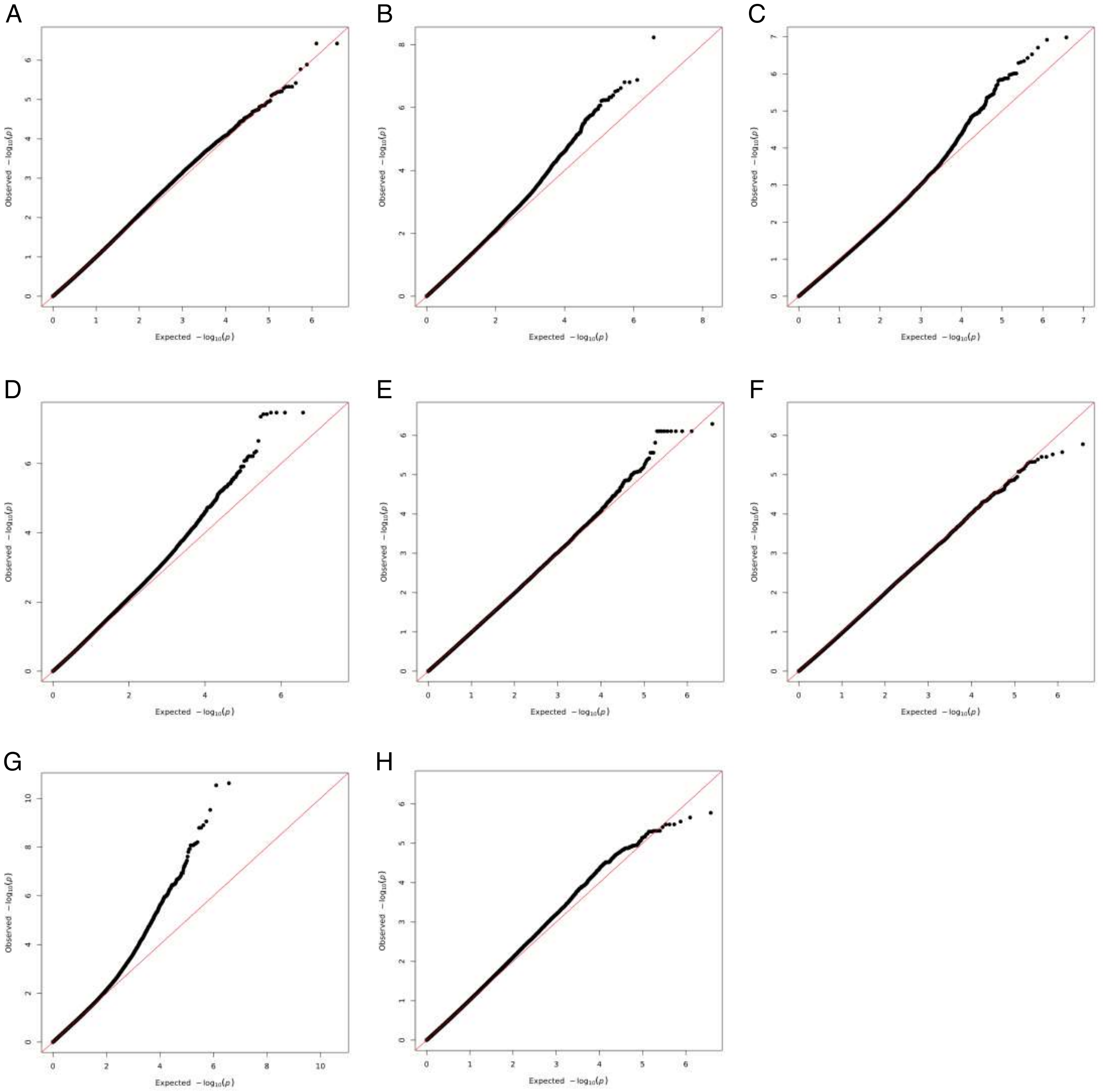


S5 Fig. Olivares et al., 2022.

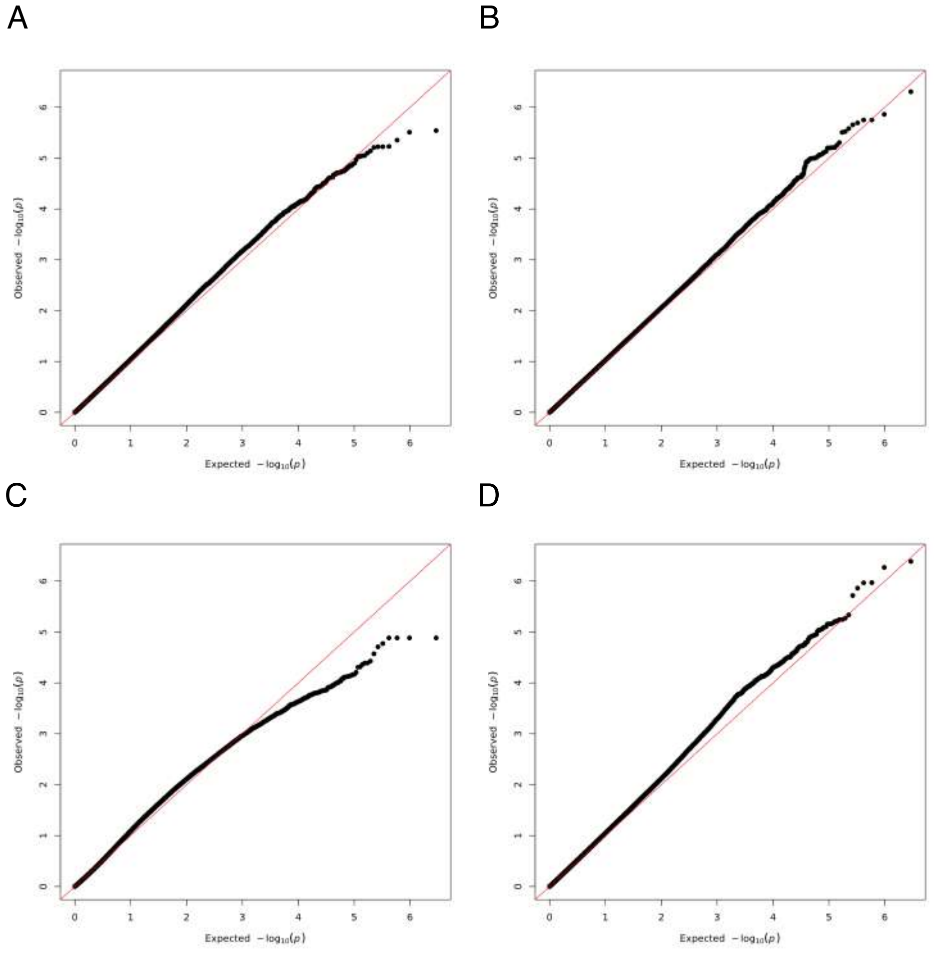


S6 Fig. Olivares et al., 2022.

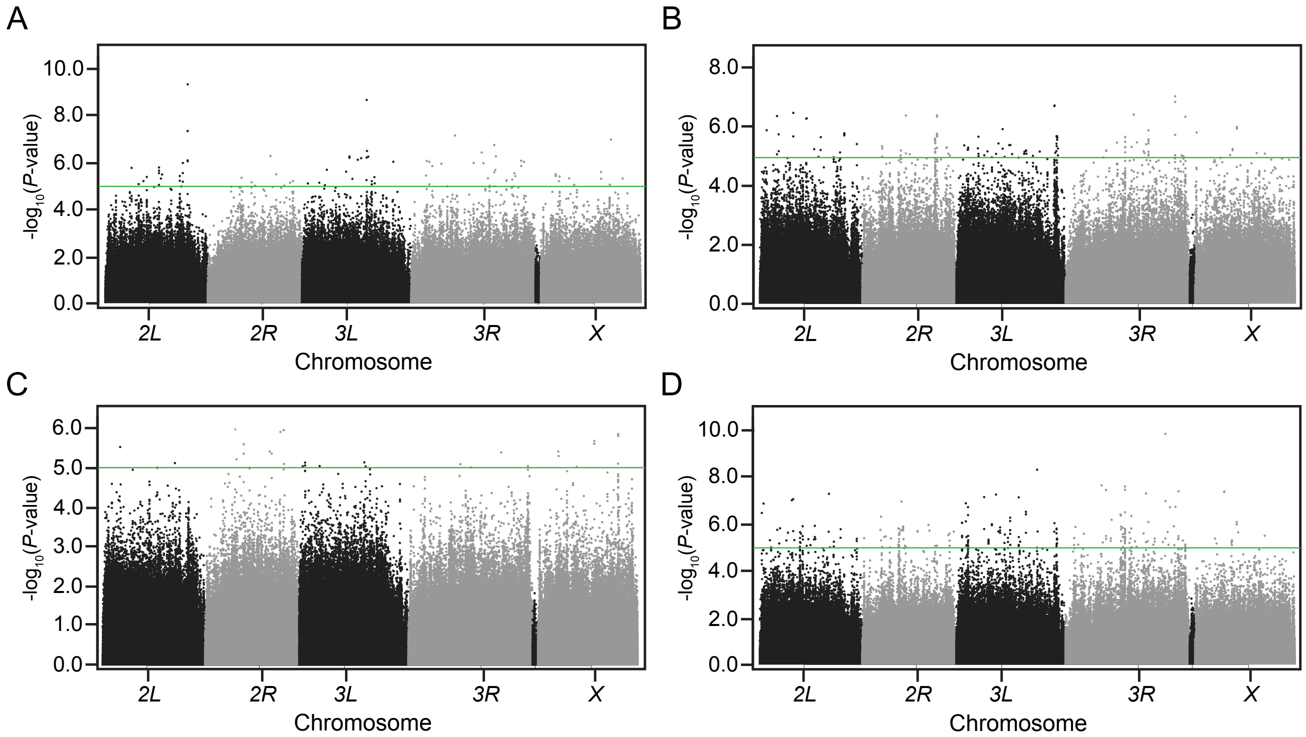


S7 Fig. Olivares et al., 2022.

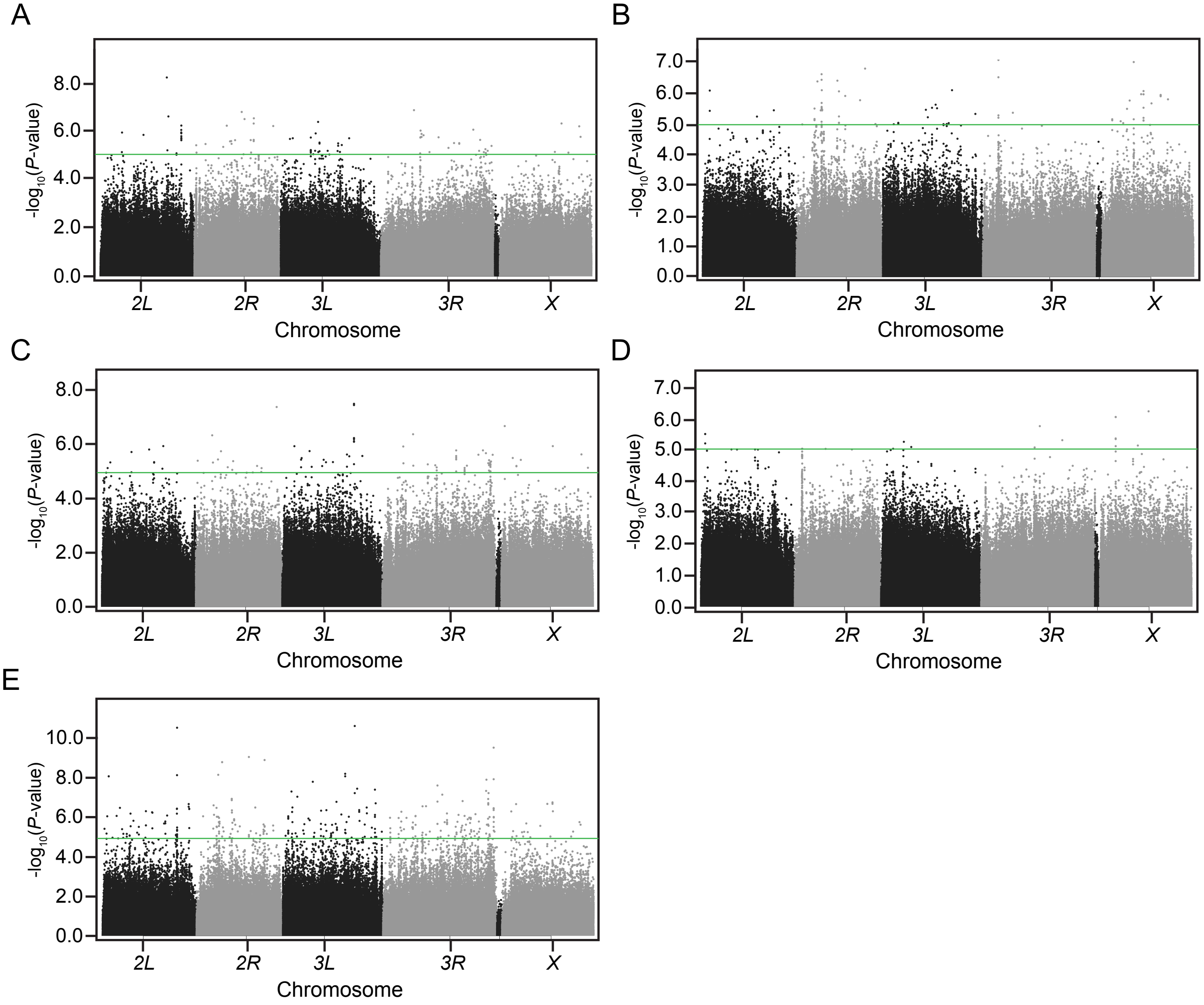


S8 Fig. Olivares et al., 2022.

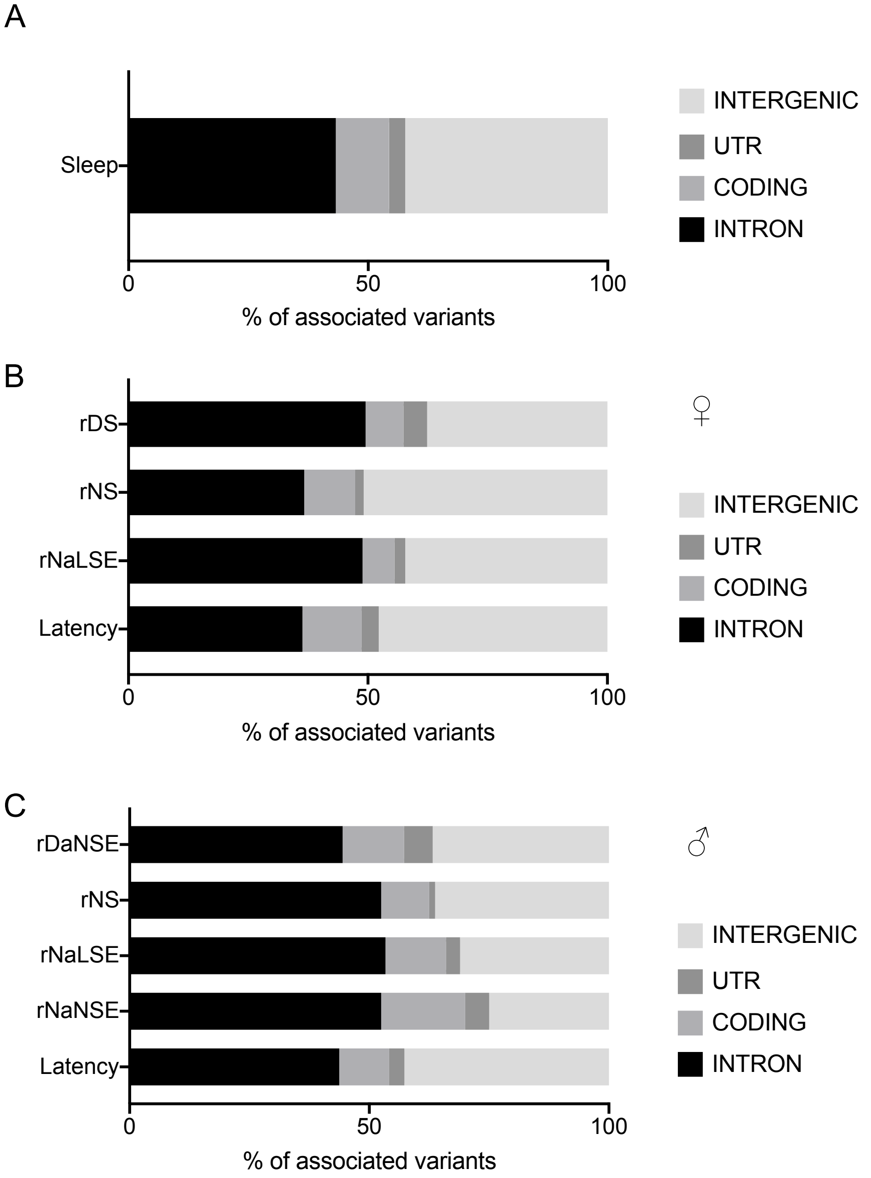


S9 Fig. Olivares et al., 2022.

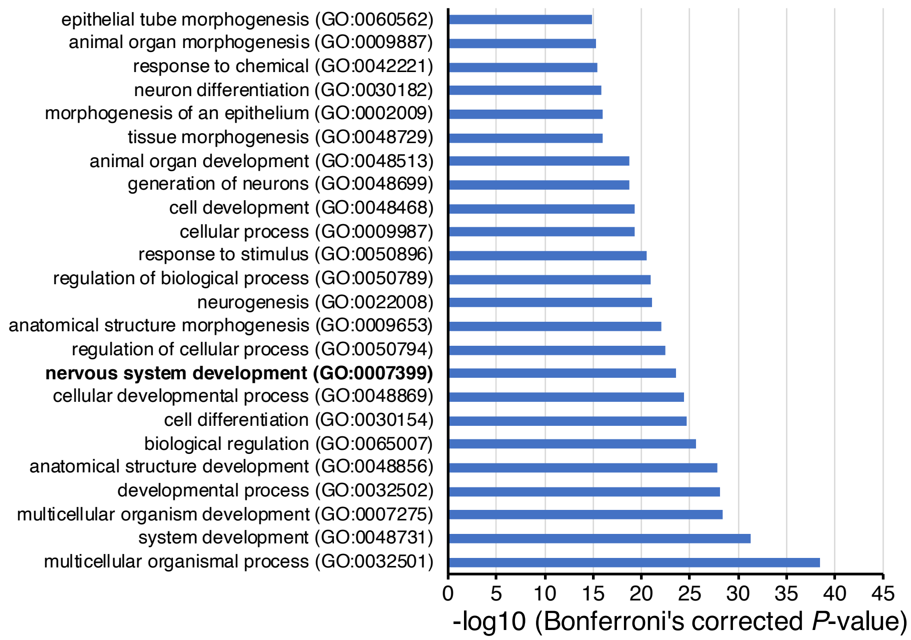


S10 Fig. Olivares et al., 2022.

A

| Fly | Human | Disease/Trait |
|---|---|---|
| *brp* | *ERC2* | Day sleep phenotypes |
| *CG11319* | *DPP10* | |
| *CG34383* | *PLEKHA5* | |
| *CG45002* | *DPP10* | |
| *Fur1* | *FURIN* | |
| *mamo* | *ZNF311* | |
| *norpA* | *PLCB4* | |
| *PsGEF* | *FRMPD2* | |
| *Ptp99A* | *PTPRG* | |
| *shn* | *HIVEP2* | |
| *shot* | *DST* | |
| *stops* | *ASB17* | |
| *tty* | *TTYH3* | |
| *beat-VI* | *F11R* | Night sleep phenotypes |
| *brp* | *ERC2* | |
| *CG11050* | *HDDC2* | |
| *CG12075* | *EPB41L4A* | |
| *JMJD5* | *KDM8* | |
| *CG30116* | *WDSUB1* | |
| *CG32373* | *FBN2* | |
| *Ca-Ma2d* | *CACNA2D3* | |
| *CG5921* | *USH1C* | |
| *CG7016* | *PSORS1C2* | |
| *CG8834* | *SLC27A6* | |
| *dpy* | *FBN2* | |
| *dsx* | *DMRT1* | |
| *PsGEF* | *USH1C* | |
| *PsGEF* | *GOPC* | |
| *PsGEF* | *LNX2* | |
| *pyd* | *TJP2* | |
| *shep* | *RBMS3* | |
| *Snmp2* | *CD36* | |
| *SoxN* | *SOX2* | |
| *vg* | *VGLL2* | |
| *CG12075* | *EPB41L3* | Sleep-related phenotypes |
| *kn* | *EBF3* | |
| *DIP-alpha* | *OPCML* | |
| *dpr10* | *OPCML* | |
| *dpr13* | *OPCML* | |
| *dpr17* | *OPCML* | |
| *dpr5* | *OPCML* | |

B

| Fly | Human | Disease/Trait |
|---|---|---|
| *PsGEF* | *PARD3B* | Hippocampal volume |
| *CG8786* | *RNF146* | Intracranial volume |
| *tau* | *MAPT* | |
| *Svil* | *SVIL* | Normalized brain volume |
| *CG8786* | *RNF146* | Subcortical brain region volumes |
| *shg* | *FAT3* | |
| *tau* | *MAPT* | |
| *kni* | *RORA* | Total ventricular volume |
| *CG12926* | *CLVS1* | Whole-brain volume |

S11 Fig. Olivares et al., 2022.

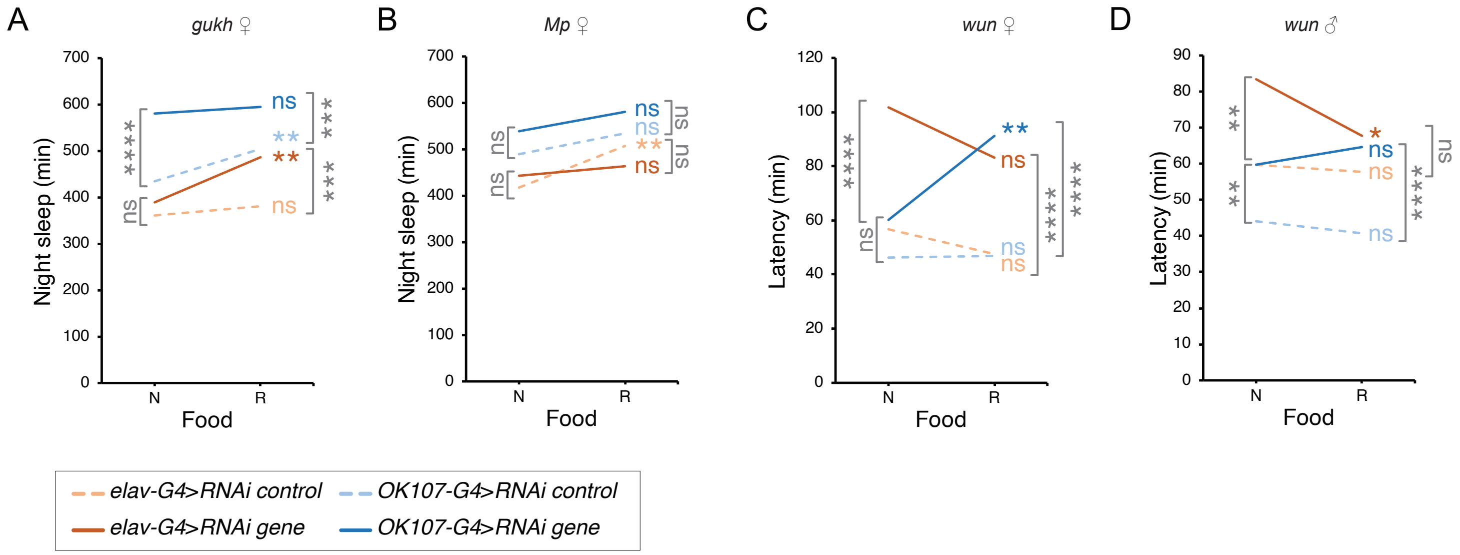


S12 Fig. Olivares et al., 2022.

A

♀

| Traits | Total sleep | | Night sleep | | Night bout number | | Night average bout length | | Day sleep | | Day bout number | | Day average bout length | | Latency | | Waking activity | | Trait of GWAS associated SNP | Number of affected Traits | |
|---|---|---|---|---|---|---|---|---|---|---|---|---|---|---|---|---|---|---|---|---|---|
| RNAi/Driver | ELAV | OK107 | ELAV | OK107 | ELAV | OK107 | ELAV | OK107 | ELAV | OK107 | ELAV | OK107 | ELAV | OK107 | ELAV | OK107 | ELAV | OK107 | | ELAV | OK107 |
| *axo* | – | ↓ | – | ↓ | – | – | – | ↓ | – | – | – | ↓ | – | ↓ | – | – | – | ↑ | Night average bout length | 0 | 6 |
| *crq* | – | ↑ | – | ↑ | – | ↑ | ↓ | – | – | ↑ | – | – | – | ↑ | – | – | – | – | Latency | 1 | 5 |
| *drongo* | – | – | – | – | – | – | ↓ | ↑ | – | – | – | – | – | ↑ | – | – | – | – | Latency | 1 | 2 |
| *Fas2* | – | ↓ | – | ↓ | – | ↑ | – | ↓ | – | – | – | – | – | ↓ | – | – | ↑ | – | Day sleep | 1 | 5 |
| *Fur1* | – | – | – | – | – | ↑ | ↓ | – | – | ↓ | – | – | – | – | – | – | – | – | | 1 | 2 |
| *gukh* | ↑ | – | ↑ | ↓ | ↓ | – | ↑ | – | – | – | – | – | – | – | ↓ | – | – | – | Night sleep | 5 | 1 |
| *LpR2* | – | – | – | – | – | ↑ | – | – | – | – | – | – | – | – | – | – | – | – | | 0 | 1 |
| *Mmp2* | – | – | – | ↓ | – | – | – | – | – | – | – | – | – | – | – | – | – | – | | 0 | 1 |
| *Mp* | – | – | ↓ | – | ↑ | ↑ | – | ↑ | – | – | – | ↓ | – | – | – | – | – | – | Night sleep | 2 | 3 |
| *mspo* | ↑ | – | ↑ | – | – | – | – | ↑ | – | – | – | ↓ | – | – | – | – | – | – | | 2 | 2 |
| *Ptp61F* | – | – | – | – | – | – | ↓ | ↑ | – | – | – | ↓ | – | – | – | ↑ | – | – | Day sleep/Night average bout length | 1 | 3 |
| *Rbfox1* | – | – | – | – | – | ↑ | ↓ | – | – | – | – | – | – | – | – | – | – | – | | 1 | 1 |
| *RpL23* | – | – | – | – | – | ↑ | ↓ | – | – | – | – | – | – | – | – | – | – | – | | 1 | 1 |
| *RpS27A* | – | – | – | – | – | – | – | – | – | – | – | – | – | ↑ | – | – | – | – | Night sleep | 0 | 1 |
| *Snmp2* | – | – | – | – | – | ↑ | ↓ | – | – | – | – | – | – | – | – | – | – | – | | 1 | 1 |
| *Ten-a* | – | ↑ | ↑ | ↑ | ↓ | – | – | ↑ | – | – | – | – | – | ↑ | – | – | ↑ | – | | 3 | 4 |
| *wun* | – | – | ↓ | – | ↑ | – | – | – | – | – | – | ↓ | ↓ | ↑ | ↓ | ↑ | – | ↑ | Latency | 4 | 4 |

B

♂

| Traits | Total sleep | | Night sleep | | Night bout number | | Night average bout length | | Day sleep | | Day bout number | | Day average bout length | | Latency | | Waking activity | | Trait of GWAS associated SNP | Number of affected Traits | |
|---|---|---|---|---|---|---|---|---|---|---|---|---|---|---|---|---|---|---|---|---|---|
| RNAi/Driver | ELAV | OK107 | ELAV | OK107 | ELAV | OK107 | ELAV | OK107 | ELAV | OK107 | ELAV | OK107 | ELAV | OK107 | ELAV | OK107 | ELAV | OK107 | | ELAV | OK107 |
| *axo* | – | – | – | – | – | – | – | – | – | – | – | – | – | – | – | – | – | – | Night average bout length | 0 | 0 |
| *crq* | – | – | – | ↓ | – | – | – | – | – | – | – | – | – | – | – | – | – | – | Latency | 0 | 1 |
| *drongo* | – | – | – | – | – | – | – | – | – | – | – | – | – | – | – | – | – | – | | 0 | 0 |
| *Fas2* | – | – | – | – | ↓ | – | ↑ | – | – | – | ↓ | – | – | – | – | – | – | – | | 3 | 0 |
| *Fur1* | – | – | – | ↓ | – | – | – | – | – | – | – | – | – | – | – | – | – | – | Latency | 0 | 1 |
| *gukh* | – | – | – | – | – | – | – | – | – | – | – | – | ↑ | – | – | – | – | – | | 1 | 0 |
| *LpR2* | – | – | – | ↓ | – | – | – | – | – | – | – | – | – | – | – | – | – | – | Latency | 0 | 1 |
| *Mmp2* | – | – | – | – | – | – | – | – | – | – | – | – | ↑ | – | – | – | – | – | Latency | 1 | 0 |
| *Mp* | – | – | – | – | – | – | – | – | – | – | – | – | – | – | – | – | – | – | | 0 | 0 |
| *mspo* | – | – | ↑ | ↓ | – | ↓ | – | ↑ | – | – | – | – | – | – | – | – | – | – | Latency | 1 | 3 |
| *Ptp61F* | – | – | – | ↓ | – | – | – | ↑ | – | – | – | – | – | – | – | – | – | – | | 0 | 2 |
| *Rbfox1* | – | – | – | ↓ | – | – | – | – | – | – | – | – | – | – | – | – | – | – | Night sleep | 0 | 1 |
| *RpL23* | – | – | – | – | – | – | – | – | – | – | – | – | – | – | – | – | – | – | Latency | 0 | 0 |
| *RpS27A* | – | – | – | ↓ | – | – | – | – | – | – | – | – | – | – | – | – | – | – | | 0 | 1 |
| *Snmp2* | – | – | – | ↓ | – | – | – | – | – | – | – | – | – | – | – | – | – | – | Latency | 0 | 1 |
| *Ten-a* | – | – | – | ↓ | – | ↓ | – | – | – | – | – | – | – | – | – | – | – | – | Night sleep | 0 | 2 |
| *wun* | – | – | – | – | – | – | – | – | ↓ | – | – | ↓ | ↓ | – | ↓ | – | – | – | Latency | 3 | 1 |

S13 Fig. Olivares et al., 2022.

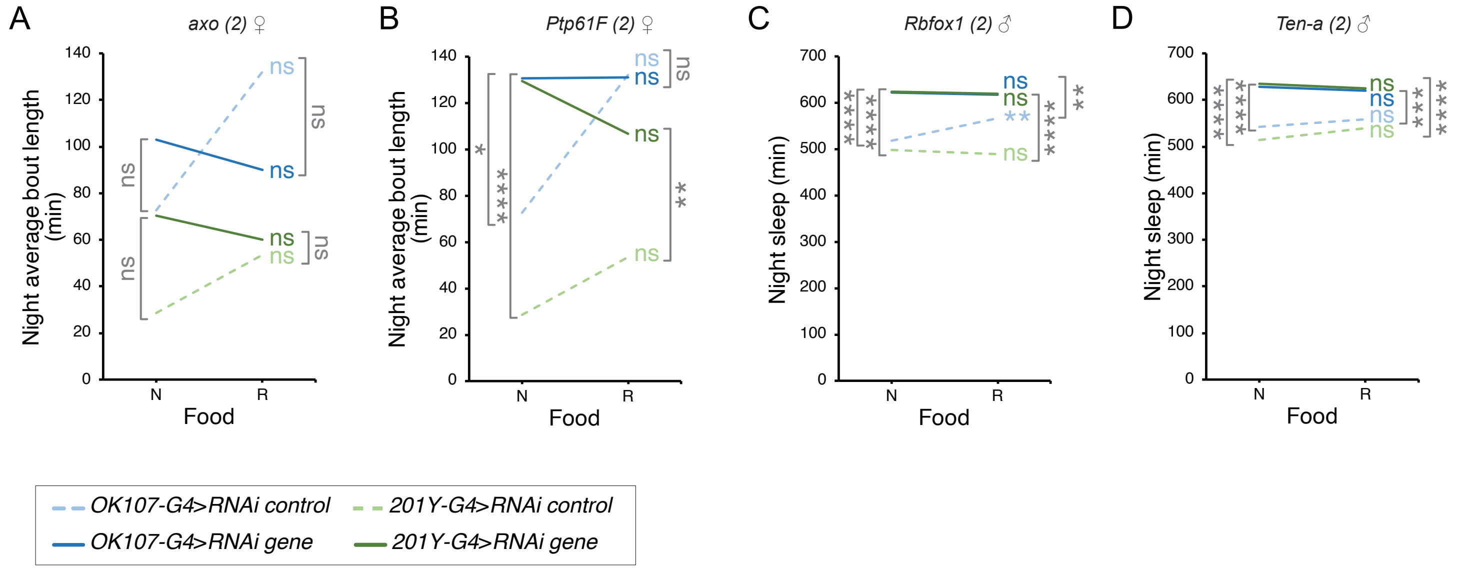


S14 Fig. Olivares et al., 2022.

A ♀

| Traits | Total Sleep | | | Night sleep | | | Night bout number | | | Night average bout length | | | Day sleep | | |
|---|---|---|---|---|---|---|---|---|---|---|---|---|---|---|---|
| RNAi/Driver | ELAV | OK107 | 201Y | ELAV | OK107 | 201Y | ELAV | OK107 | 201Y | ELAV | OK107 | 201Y | ELAV | OK107 | 201Y |
| *axo* | – | ↓ | – | – | ↓ | – | – | – | – | – | ↓ | – | – | – | – |
| *axo (2)* | ND | – | – | ND | – | ↓ | ND | ↑ | ↑ | ND | – | – | ND | – | – |
| *Ptp61F* | – | – | – | – | – | – | – | – | – | ↓ | ↑ | – | – | – | – |
| *Ptp61F (2)* | ND | – | – | ND | – | ↓ | ND | ↑ | ↑ | ND | – | – | ND | – | – |
| *Rbfox1* | – | – | – | – | – | ↑ | – | ↑ | ↓ | ↓ | – | – | – | – | – |
| *Rbfox1 (2)* | ND | – | – | ND | – | – | ND | ↑ | – | ND | – | – | ND | – | – |
| *Ten-a* | – | ↑ | ↑ | ↑ | ↑ | ↑ | ↓ | – | ↓ | – | ↑ | – | – | – | – |
| *Ten-a (2)* | ND | – | – | ND | – | ↓ | ND | ↑ | ↑ | ND | – | – | ND | – | ↓ |

| Traits | Day bout number | | | Day average bout length | | | Latency | | | Waking activity | | | Trait of GWAS associated SNP | Number of affected Traits | | |
|---|---|---|---|---|---|---|---|---|---|---|---|---|---|---|---|---|
| RNAi/Driver | ELAV | OK107 | 201Y | ELAV | OK107 | 201Y | ELAV | OK107 | 201Y | ELAV | OK107 | 201Y | | ELAV | OK107 | 201Y |
| *axo* | – | ↓ | – | – | ↓ | – | – | – | – | – | ↑ | – | | 0 | 6 | 0 |
| *axo (2)* | ND | – | – | ND | – | – | ND | – | – | ND | – | ↓↓ | | ND | 1 | 3 |
| *Ptp61F* | – | ↓ | – | – | – | – | – | ↑ | – | – | – | – | Day sleep / Night average bout length | 1 | 3 | 0 |
| *Ptp61F (2)* | ND | – | – | ND | – | | ND | – | – | ND | – | ↓ | | ND | 1 | 3 |
| *Rbfox1* | – | – | – | – | – | – | – | – | – | – | – | – | | 1 | 1 | 2 |
| *Rbfox1 (2)* | ND | – | – | ND | – | – | ND | – | – | ND | – | – | | ND | 1 | 0 |
| *Ten-a* | – | – | ↓ | – | ↑ | – | – | – | ↓ | ↑ | – | – | | 3 | 4 | 5 |
| *Ten-a (2)* | ND | – | – | ND | – | – | ND | – | – | ND | – | ↓ | | ND | 1 | 4 |

B ♂

| Traits | Total sleep | | | Night sleep | | | Night bout number | | | Night average bout length | | | Day sleep | | |
|---|---|---|---|---|---|---|---|---|---|---|---|---|---|---|---|
| RNAi/Driver | ELAV | OK107 | 201Y | ELAV | OK107 | 201Y | ELAV | OK107 | 201Y | ELAV | OK107 | 201Y | ELAV | OK107 | 201Y |
| *axo* | – | – | – | – | – | – | – | – | – | – | – | – | – | – | – |
| *axo (2)* | ND | – | – | ND | – | – | ND | – | ↑ | ND | – | – | ND | – | – |
| *Ptp61F* | – | – | – | – | ↓ | – | – | – | – | – | ↑ | – | – | – | – |
| *Ptp61F (2)* | ND | – | – | ND | – | – | ND | – | ↑ | ND | – | – | ND | – | – |
| *Rbfox1* | – | – | – | – | ↓ | – | – | – | – | – | – | – | – | – | – |
| *Rbfox1 (2)* | ND | – | – | ND | ↓ | – | ND | – | – | ND | – | – | ND | – | – |
| *Ten-a* | – | – | – | – | ↓ | – | – | ↓ | – | – | – | – | – | – | – |
| *Ten-a (2)* | ND | – | – | ND | – | – | ND | – | ↑ | ND | – | – | ND | – | – |

| Traits | Day bout number | | | Day average bout length | | | Latency | | | Waking activity | | | Trait of GWAS associated SNP | Number of affected Traits | | |
|---|---|---|---|---|---|---|---|---|---|---|---|---|---|---|---|---|
| RNAi/Driver | ELAV | OK107 | 201Y | ELAV | OK107 | 201Y | ELAV | OK107 | 201Y | ELAV | OK107 | 201Y | | ELAV | OK107 | 201Y |
| *axo* | – | – | – | – | – | – | – | – | – | – | – | – | | 0 | 0 | 0 |
| *axo (2)* | ND | – | – | ND | – | – | ND | – | – | ND | – | – | | ND | 0 | 1 |
| *Ptp61F* | – | – | – | – | – | – | – | – | – | – | – | ↓ | | 0 | 2 | 1 |
| *Ptp61F (2)* | ND | – | – | ND | – | – | ND | – | – | ND | – | – | | ND | 0 | 1 |
| *Rbfox1* | – | – | – | – | – | – | – | – | – | – | – | – | Night sleep | 0 | 1 | 0 |
| *Rbfox1 (2)* | ND | – | – | ND | – | – | ND | – | – | ND | – | – | | ND | 1 | 0 |
| *Ten-a* | – | – | ↓ | – | – | – | – | – | – | – | – | – | Night sleep | 0 | 2 | 1 |
| *Ten-a (2)* | ND | – | – | ND | – | – | ND | – | – | ND | – | – | | ND | 0 | 1 |

S15 Fig. Olivares et al., 2022.

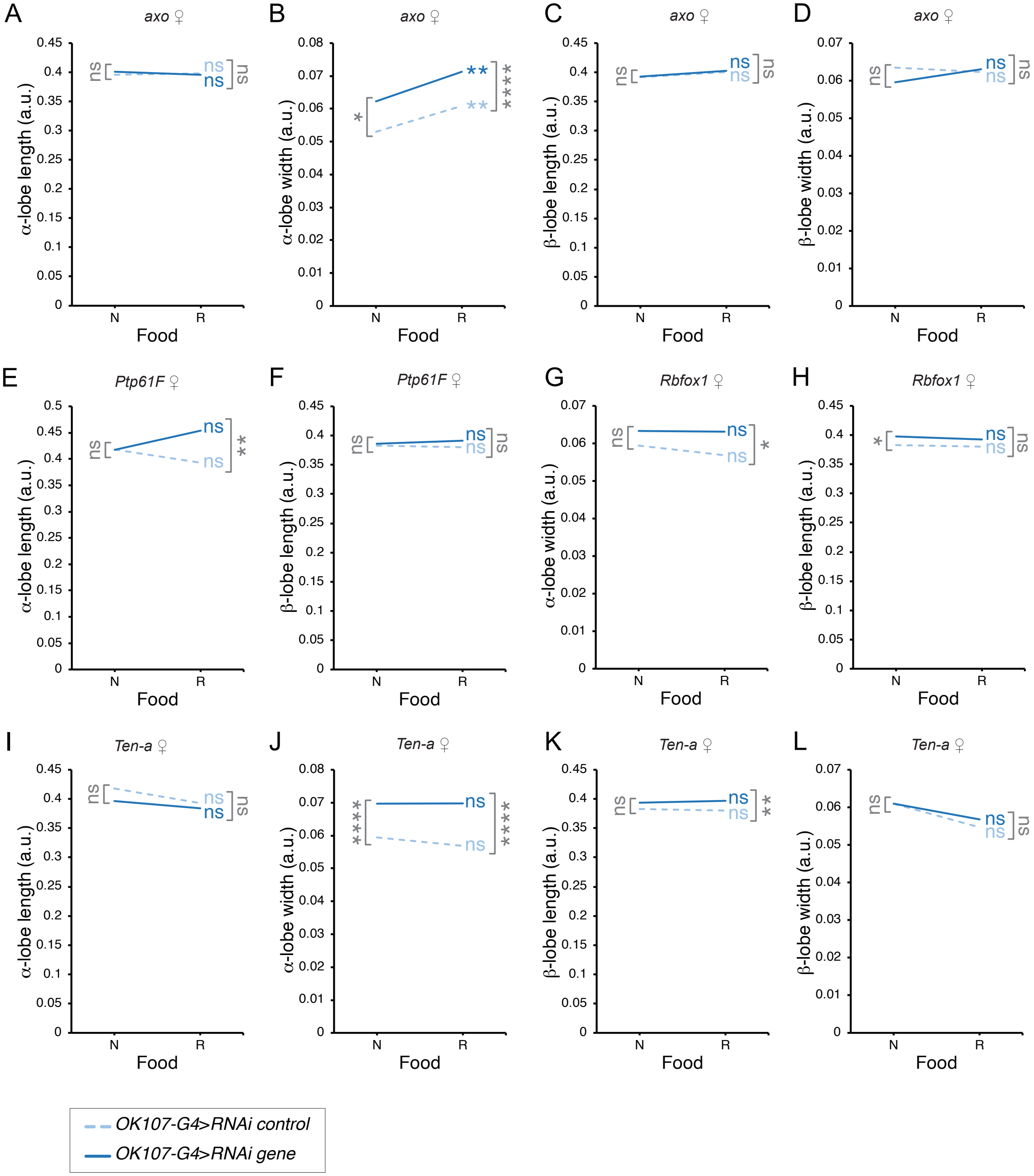


S16 Fig. Olivares et al., 2022.

| Traits | α-lobe length | α-lobe width | β-lobe length | β-lobe width |
|---|---|---|---|---|
| RNAi/Driver | *OK107* | *OK107* | *OK107* | *OK107* |
| *axo* | – | – | – | – |
| *Ptp61F* | – | ↑ | – | ↑ |
| *Rbfox1* | ↓ | – | – | ↓ |
| *Ten-a* | – | – | – | – |

| Number of affected Traits |
|---|
| *OK107* |
| 0 |
| 2 |
| 2 |
| 0 |

♀

S17 Fig. Olivares et al., 2022.
